# Combining functional annotation and multi-trait fine-mapping methods improves fine-mapping resolution at glycaemic trait loci

**DOI:** 10.1101/2024.11.13.623384

**Authors:** Jana Soenksen, Ji Chen, Arushi Varshney, Susan Martin, MAGIC, Stephen C. J. Parker, Andrew P. Morris, Jennifer L. Asimit, Inês Barroso

## Abstract

The Meta-Analysis of Glucose and Insulin-related traits Consortium (MAGIC) identified 242 loci associated with glycaemic traits fasting insulin (FI), fasting glucose (FG), 2h-Glucose (2hGlu), and glycated haemoglobin (HbA1c). However, for the majority, the causal variant(s) remain(s) unknown. Modelling multiple traits and integrating functional annotations have each been shown to improve fine-mapping resolution. Here, we aimed to determine whether combining these techniques would further improve fine-mapping resolution.

Using single-trait fine-mapping results from FINEMAP as input, we performed multi-trait fine-mapping with flashfm at 50 loci significantly associated with more than one glycaemic trait. We used fGWAS to build models of enriched annotations by considering 32 cell-type specific and 28 static annotations. We used the prior probabilities from these models to perform annotation informed fine-mapping with both FINEMAP (single-trait) and flashfm (multi-trait).

Multi-trait fine-mapping of 106 locus-trait associations significantly (p=1.23 x 10^-17^) reduced the median size of the credible sets accounting for 99% of the posterior probability of being causal (99CS) to 21.5 variants compared to the 60.5 variants in single-trait fine-mapping. Annotation informed single-trait fine-mapping of 211 locus-trait associations reduced (p=4.24x10^-12^) the median 99CS size from 72 in agnostic single-trait fine-mapping to 52 variants. Annotation informed multi-trait fine-mapping of 110 locus-trait associations led to a further significant (p=2.69x10^-18^) decrease in median 99CS size to 14.5 variants compared to 51.0 in annotation informed single-trait fine-mapping.

In conclusion, we found that multi-trait and annotation informed fine-mapping can help to further narrow down likely causal variants at glycaemic trait loci, both separately and when combined.

**Author Summary:** Large-scale studies such as the Meta-analysis of glucose and insulin-related traits consortium (MAGIC) identified regions in our DNA which affect glycaemic measures related to type 2 diabetes, such as glucose and insulin levels measured after fasting overnight. However, these regions contain many DNA changes within them that are close to each other and that are inherited together. This makes identifying the DNA change responsible for the effect on glycaemic measures challenging. Fine-mapping is a statistical approach that helps to narrow down the list of DNA changes that are likely to be responsible for the association with glycaemic measures, referred to as causal DNA changes. Here we used multiple types of fine-mapping approaches together to combine their advantages. We showed that both combining data from multiple related glycaemic measures as well as including information about the likely function of DNA changes helps to narrow down the number of likely causal DNA changes. Combining both approaches led to the biggest improvement in the list of likely causal DNA changes. While this study focuses on glycaemic measures, the approaches can be applied to other measures used to monitor human health, for example blood pressure or cholesterol levels. This study highlights the benefits of combining multiple approaches to find likely causal DNA changes.

## Introduction

Genetic studies of fasting and post-prandial measures of insulin and glucose, and glycated haemoglobin (HbA1c) levels have provided insights into pathways related to glucose homeostasis and diabetes pathophysiology [1–3]. The largest multi-ancestry study from MAGIC (Meta-Analysis of Glucose and Insulin-related traits Consortium) identified 242 loci associated with glycaemic traits fasting insulin (FI), fasting glucose (FG), 2-hour glucose (2hGlu) and HbA1c [1]. The authors demonstrated that multi-ancestry fine-mapping improved resolution compared to single-ancestry fine-mapping when comparing results with equivalent sample sizes [1]. However, fine-mapping could not resolve the majority of the loci to a single causal variant. Recently, fine-mapping methods that leverage information across multiple traits have been developed [4]. In addition, methods that leverage functional annotation have also been shown to improve fine-mapping resolution [5, 6].

Here, we used association results from Chen et al [1] to investigate whether multi-trait fine-mapping would improve fine-mapping resolution by leveraging information across multiple glycaemic traits. We compared the results to single-trait fine-mapping performed in the European (EUR)-like group and to multi-ancestry fine-mapping from Chen et al [1]. In addition, taking advantage of the functional annotations known to be enriched for each of the glycaemic traits [1], we explored whether the addition of this information to both single-trait and multi-trait fine-mapping approaches would help to further improve fine-mapping resolution, and identify the underlying causal variants. We showed that these approaches successfully improved fine-mapping resolution at glycaemic trait loci.

## Results

### Identification of loci for fine-mapping

Of the 234 loci fine-mapped by Chen et al [1], we focused on 155 loci that were associated (p-value<5x10^-8^) with at least one of the glycaemic traits, FI, FG, HbA1c and 2hGlu, in EUR-like discovery data (**Supplementary table 1 & Supplementary Figure 1**). Of these, two loci near *GCK* and *G6PC2* were excluded because previous fine-mapping had demonstrated multiple independent variants at each locus, and their roles in glucose homeostasis are well-established [7–9]. We noted that three loci associated with FG, 2hGlu and FI overlapped, and given the possibility of shared common causal variant(s), we merged them into a single locus and repeated single-trait fine-mapping for this new larger locus (**Methods**). We carried out annotation-informed fine-mapping at the resulting 151 loci. A subset of 50 loci was associated with a second glycaemic trait with p-value<10^-6^ in EUR-like fine-mapping datasets (**Methods**). We carried out agnostic and annotation informed fine-mapping at these 50 loci (**Supplementary table 1**).

There is no single best metric to compare performance of different fine-mapping approaches and as we do not have the ground truth, we focused on the size of the 99 % credible set (99% CS), and the marginal posterior probability (MPP) of causality of individual variants (**Methods**). Although neither of these metrics guarantee that the causal variant has been captured, smaller 99% CS and higher MPP values provide increased confidence in the fine-mapping results. These metrics are commonly used to assess performance in fine-mapping analyses, as they prioritise smaller sets of variants with higher likelihood of being causal for downstream functional studies. For these reasons, we have chosen to focus on these metrics.

### Multi-trait fine-mapping improves resolution compared to single-trait fine-mapping

We used the single-trait fine-mapping results at 50 loci (110 locus-trait associations), as input for multi-trait fine-mapping. All but one locus (4-locus trait associations) were successfully run, leading to 106 locus-trait associations with agnostic multi-trait fine-mapping results (**Methods**). When comparing the overall performance of the multi-trait to single-trait fine-mapping, the median size of the 99% CS was reduced from 60.5 (lower quartile = 18.3, upper quartile = 314) in single-trait to 21.5 (lower quartile = 7.25, upper quartile = 80.0) variants in multi-trait (p=1.23 x 10^-17^, **Figure 1 A & C**). All 99% CS sizes either stayed the same or were reduced in multi-trait fine-mapping. Furthermore, the median MPP of the variant with the highest MPP (top variant) significantly increased from 0.303 (lower quartile = 0.166, upper quartile = 0.563) to 0.410 (lower quartile = 0.236, upper quartile = 0.752) in multi-trait fine-mapping (p=2.65x10^-7^, **Figure 1 B**). Overall, multi-trait fine-mapping led to more locus-trait associations with smaller 99% CS sizes compared to single-trait fine-mapping (**Figure 1 C**). For example, the number of locus trait-associations with 2-5 variants in the 99% CS increased from 5/106 in single-trait fine-mapping to 18/106 in multi-trait fine-mapping (**Figure 1 C**).

**Figure 1.**
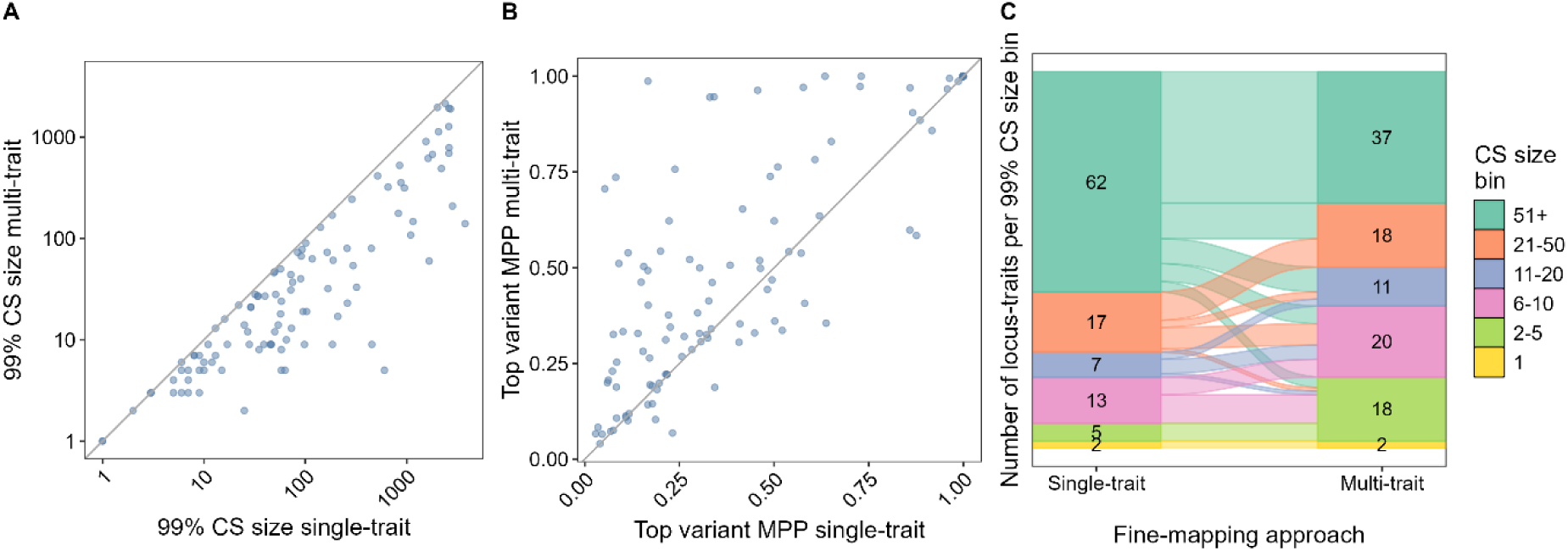
Comparison of multi-trait and single-trait fine-mapping results A&B: Each dot represents the respective results at a locus-trait association according to the single-trait fine-mapping approach FINEMAP v1.4 (x-axis) compared to the multi-trait approach Flashfm (y-axis), both are single-ancestry analysis performed in EUR-like populations. The grey line represents the line of equality. Multi-trait results were obtained using the single-trait results as input. **A**: Number of SNPs in the 99% CS. **B**: Marginal posterior probability (MPP) of the variant with the highest MPP at each locus-trait association. **C**: Comparison of the number of locus-trait associations in each 99% CS size bins between different methods (indicated by number in bar chart) and illustrates how the bins of locus-traits change between methods.

### Overall fine-mapping performance is similar for multi-ancestry and multi-trait fine-mapping

Both multi-trait and multi-ancestry fine-mapping can improve fine-mapping resolution compared to single-trait single-ancestry fine-mapping. However, depending on the datasets available multi-ancestry fine-mapping might not be possible. Furthermore, at some loci multi-ancestry fine-mapping performs worse than single-ancestry fine-mapping likely due to effect size heterogeneity between different populations [1]. We compared the resolution of multi-trait fine-mapping with that of multi-ancestry fine-mapping at the subset of 93 locus-trait associations which were also significantly associated in multi-ancestry data (log10BF>6, **Figure 2**). The median 99% CS size of multi-ancestry fine-mapping was 24.0 compared to 21.0 in multi-trait fine-mapping (p=0.540, **Figure 2 A & C**). The 99% CS size in multi-trait analysis was reduced in 40 of 93 locus-trait associations (45.4%) compared to multi-ancestry and was the same size in another 10 locus-trait associations. This highlights that in some cases fine-mapping resolution improved compared to multi-ancestry fine-mapping even though the overall difference was not significant and the sample size was smaller in the multi-trait analysis (**Figure 2 A**). The median MPP of the top variant was 0.522 in multi-ancestry and 0.407 in multi-trait fine-mapping (p=0.118, **Figure 2 B & D**). Overall, these results show no significant difference in performance of multi-trait and multi-ancestry fine-mapping.

The best performing fine-mapping approach varied between locus-trait associations. To understand why multi-trait fine-mapping might outperform multi-ancestry fine-mapping at 40 locus-trait associations, we compared the results between single-trait single-ancestry and multi-ancestry fine-mapping (**Supplementary Figure 2**). Of these locus-trait associations with improved multi-trait fine-mapping, 17/40 had performed worse with multi-ancestry fine-mapping compared to single-ancestry single-trait fine-mapping. However, at 23 locus-trait associations, the multi-trait fine-mapping improved over the already improved multi-ancestry fine-mapping. This showed that multi-trait fine-mapping can outperform multi-ancestry fine-mapping, and suggests that performing multi-trait fine-mapping may be helpful to refine association signals even if multi-ancestry fine-mapping has already been conducted.

**Figure 2.**
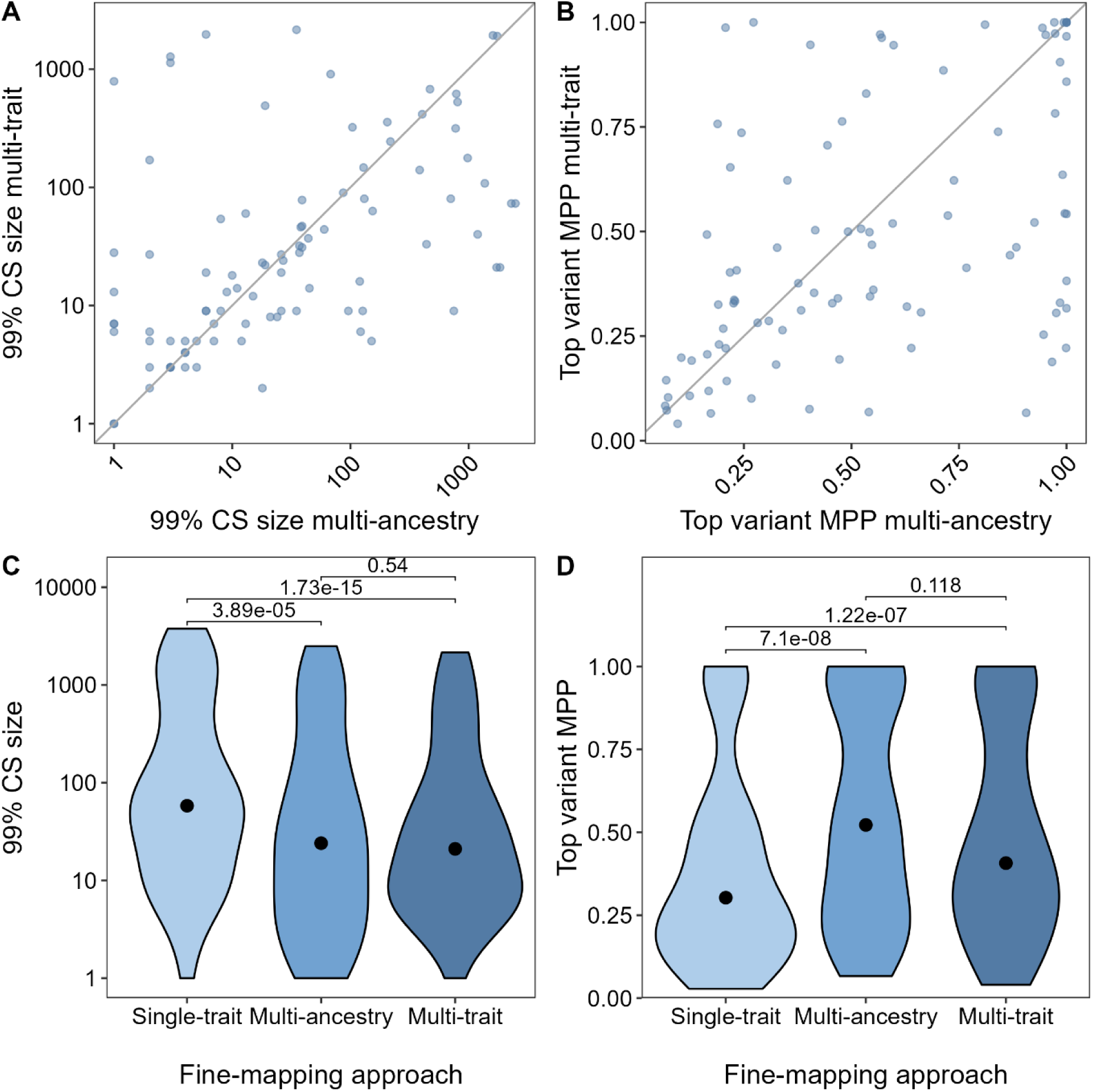
Comparison of agnostic fine-mapping approaches. Multi-trait fine-mapping was carried out with flashfm and using single-trait single ancestry fine-mapping results from EUR-like data obtained with FINEMAP v1.4 as input. Multi-ancestry fine-mapping was carried out using FINEMAP v1.1 in a single trait **A&B**: Each dot represents the results at a locus-trait association according to multi-ancestry (x-axis) compared to multi-trait approach (y-axis). The grey line represents the line of equality. The multi-trait results were obtained using the single-trait results from EUR-like as input. **C&D**: Includes only locus-trait associations that were analysed with all three approaches. The dot indicates the median, p-values were calculated with a paired two sided Wilcoxon. **A&C**: Compares the number of SNPs in the 99% credible set (99% CS). **B&D**: The marginal posterior probability (MPP) of the variant with the highest MPP of being causal.

Annotation-informed multi-trait fine-mapping improves fine-mapping resolution compared to annotation-agnostic and annotation-informed single-trait fine-mapping So far, all fine-mapping approaches were annotation-agnostic and not informed by any known function of variants. We next sought to explore if using annotation-informed fine-mapping would improve resolution compared to annotation-agnostic single-trait fine-mapping (**Supplementary Figure 3**) and if this could further be improved by conducting annotation-informed multi-trait fine-mapping (**Supplementary Figure 3**).

Starting with the 28 static annotations and 32 cell-type specific stretch enhancers previously analysed by Chen et al [1], we built models of enriched annotations to calculate priors for annotation-informed fine-mapping (**Methods**). The resulting models contained between two (2hGlu) and six annotations (FG and HbA1c, **Supplementary Figure 4**). Static annotations such as coding or conserved regions were included in the final model of multiple glycaemic traits (FG, 2hGlu and HbA1c). FG and 2hGlu results were enriched for islet stretch enhancers, while FI was enriched for adipose-specific stretch enhancers.

After performing annotation-informed fine-mapping using these priors we compared the results with agnostic fine-mapping results at 211 locus-trait associations in EUR-like, including all loci associated with only a single trait (**Methods**). The median 99% CS size decreased significantly from 72.0 (lower quartile = 21.5, upper quartile = 458) variants in agnostic single-trait fine-mapping to 52.0 (lower quartile = 15.5, upper quartile = 319) after adding annotations (p=4.24x10^-12^, **Supplementary Figure 3 A**). Furthermore, there was a significant increase in the median MPP of the top variant from 0.277 in agnostic fine-mapping to 0.418 after adding annotation (p=6.83x10^-15^, **Supplementary Figure 3 C**). Not all locus-trait associations were improved to the same extent. For example, at the *ZBED3-AS1* locus, adding annotations to single trait fine-mapping decreased the 99% CS size more strongly in FG (from 63 to 8 variants) compared to HbA1c (from 15 to 10 variants). At 36 locus-trait associations, adding annotations increased the 99% CS size (**Supplementary Figure 3 A**). However, at a subset of 13 locus-trait associations the MPP of the top variant increased which might suggest the overall fine-mapping did not perform worse at these loci.

We showed that multi-trait fine-mapping and annotation-informed fine-mapping both significantly improved the resolution of the relevant loci. We next combined these two approaches and used annotation-informed fine-mapping on the same subset of 110 locus-trait associations that we had fine-mapped with agnostic multi-trait fine-mapping (**Methods**). The median number of SNPs in the 99% CS decreased from 51.0 (lower quartile = 11.3, upper quartile = 305) variants in annotation-informed single-trait fine-mapping to 14.5 (lower quartile = 5.00, upper quartile = 73.5) in annotation-informed multi-trait fine-mapping (p=2.69 x 10^-18^, **Supplementary Figure 3 B**). The 99% CS size either stayed the same or decreased for all locus-trait associations in multi-trait fine-mapping compared to single-trait fine-mapping. The median MPP of the top variant increased significantly from 0.444 in single-trait to 0.706 in multi-trait fine-mapping (p = 1.47 x 10^-15^, **Figure 3 A & Supplementary Figure 3 D**).

Of the 110 locus-trait associations, 2 resolved to a single variant in the 99% CS in agnostic single-trait fine-mapping. This improved to 3 in annotation-informed single-trait fine-mapping and 8 in annotation-informed multi-trait fine-mapping (**Figure 3 B**). Furthermore, there were 28 locus-trait associations with 2-5 variants in the 99% CS with annotation-informed multi-trait fine-mapping, an increase from 6 in the agnostic single-trait fine-mapping results (**Figure 3 B**).

Overall, when comparing fine-mapping results from single-trait agnostic, to single-trait and multi-trait annotation-informed methods there was a gradual improvement of results (**Figure 3 A & B**). However, the 99% CS of some locus-trait associations were not decreased by adding functional annotation. Therefore, there isn’t a single best performing method and using multiple approaches is required to find the smallest 99% CS.

**Figure 3.**
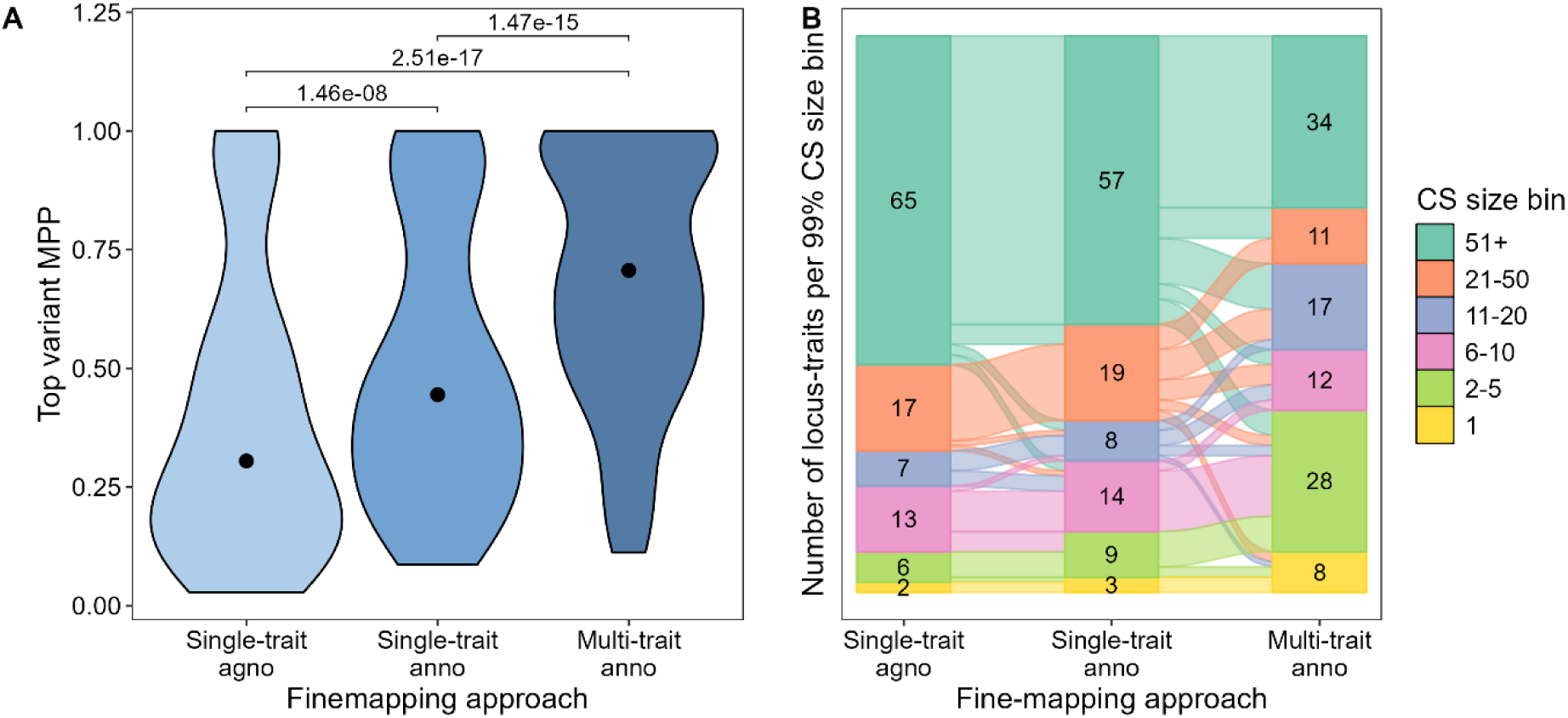
Annotation-informed fine-mapping approaches. Single trait fine-mapping was carried out with FINEMAP v1.4. Annotation (anno) informed single-trait fine-mapping was carried out using priors calculated with fGWAS. Multi-trait annotation-informed fine-mapping was carried out with flashfm using single-trait annotation-informed results as input. **A** The marginal posterior probability (MPP) of the variant with the highest MPP of being causal. The dot indicates the median, p-values were calculated with a paired two-sided Wilcoxon. **B:** Compares the number of locus-trait associations in 99% credible set (99% CS) size bins between different methods (indicated by number in bar chart) and illustrates how the bins of locus-traits change between methods

## Discussion

In this study, we aimed to investigate multiple fine-mapping approaches, including multi-trait and functionally informed fine-mapping, and assess their impact on fine-mapping resolution at glycaemic trait loci. Firstly, we applied multi-trait fine-mapping at loci associated with more than one trait. This significantly improved fine-mapping resolution and led to 99% CS sizes either staying the same or decreasing compared to fine-mapping in EUR-like only data. Multi-trait fine-mapping resolution was not significantly different to results obtained from multi-ancestry fine-mapping. Secondly, we used annotation-informed single-trait and multi-trait fine-mapping which led to an overall significant decrease of 99% CS sizes.

There was no significant difference between the performance of multi-trait and multi-ancestry fine-mapping approaches. This highlights that multi-trait fine-mapping can be a good alternative to multi-ancestry fine-mapping if data from diverse ancestries is not available. Furthermore, performing both alongside each other can improve fine-mapping for individual locus-trait associations, especially for loci that performed worse in multi-ancestry single-trait fine-mapping compared to single-ancestry single-trait fine-mapping. Multi-ancestry fine-mapping might perform worse due to effect size heterogeneity between ancestries, due to differences in the linkage disequilibrium (LD), or interactions between genes and environment. However, one limitation of our study was that the sample sizes in the multi-ancestry fine-mapping and the multi-trait fine-mapping were not matched so some of the improvements in multi-ancestry fine-mapping could additionally be driven by increased power due to overall larger sample size.

Adding annotations to single-trait fine-mapping significantly improved the overall fine-mapping resolution, but in some cases adding annotations increased the CS size. This demonstrates the value of performing fine-mapping unbiased by current functional knowledge of the genome. If the true causal variant does not map to one of the annotations included in the model or its function is not known at all, adding annotation information might worsen the performance of fine-mapping at this locus as it might give higher MPPs to non-causal but annotated variants with a higher prior probability. This explains some of the increased 99% CS sizes after adding annotations.

Relatedly, at the *ZBED3-AS1* locus associated with HbA1c and FG, fine-mapping produced smaller 99% CS sizes for HbA1c than for FG when comparing the corresponding agnostic fine-mapping approaches. Conversely, the annotation-informed fine-mapping methods led to smaller 99% CS sizes for FG than HbA1c. A potential explanation is that the fGWAS model for FG includes introns as enriched while the HbA1c model does not. All three variants in the 99% CS of FG are intronic variants of *ZBED3*-*AS1* and these three variants are also included in the 99% CS of HbA1c. Since the final HbA1c enrichment model does not contain intronic variants these had a lower prior than those in FG. The differences between the annotation models likely affected the extent of the decrease in 99% CS. Functional evidence at this locus supports the causal variant at this locus being the intronic variant rs7732130; Miguel-Escalada et al [10] identified rs7732130 as being part of an enhancer hub using islet promoter capture Hi-C interaction data. CRISPRa and CRISPRi perturbation of one of these variants, rs7732130 (MPP=0.873), led to gene expression changes of *ZBED3, snoRA47, ZBED3-AS1, PDE8B* and *WDR41* [10]. *ZBED3-AS1* has been shown to regulate *ZBED3* expression which in turn affects Wnt/beta catenin signalling pathway and adipocyte differentiation [11]. *ZBED3* also supressed insulin induced phosphorylation of the insulin receptor and Akt [12]. These pathways can all have an effect on glucose metabolism which is reflected in elevated glucose levels in a mouse knock-out model of *Zbed3* [13]. Given the functional evidence and mechanism linking the 99% CS variant rs7732130 to glucose metabolism via *ZBED3* expression, annotation-informed fine-mapping likely prioritised the right variant for both FG and HbA1c associations at the locus.

In conclusion, for loci associated with multiple traits, conducting multi-trait fine-mapping improved resolution with or without adding annotations. If multi-ancestry fine-mapping is not available or did not improve fine-mapping at a locus, carrying out multi-trait fine-mapping can be a valuable alternative. Finally, adding annotations improved overall fine-mapping resolution but if the causal variant is not part of one of the annotations in the final model it can lead to worse results. Therefore, it is important to consider both agnostic and annotation-informed approaches to find the most likely causal variant.

## Methods

### Genetic and genomic functional annotation datasets used

We used trans-ancestry (here called multi-ancestry) and European ancestry (here called EUR-like) meta-analysis summary statistics from MAGIC (http://magicinvestigators.org/downloads/). As per Chen et al [1], cohorts that were only genotyped in the Metabochip were excluded from fine-mapping analysis and FG, 2hGlu and FI were adjusted for BMI (**Supplementary table 2**).

For analysis of enriched functional annotations, we used static annotations from Finucane et al [14] and cell-type specific annotations, i.e. stretch enhancers from Varshney et al [15] as described by Chen et al [1].

### Locus selection for multi-trait fine-mapping

Loci were selected for multi-trait fine-mapping if they were genome-wide significantly associated in the discovery data set (p<5x10^-8^, including Metabochip data, **Supplementary table 1**) with at least one of the glycaemic traits, FI, FG, 2hGlu or HbA1c in EUR-like individuals (210 locus-trait associations), based on results from MAGIC [1]. We excluded from fine-mapping loci that were not genome-wide significant in multi-ancestry meta-analysis or if they mapped to the X chromosome. In agreement with recommendations from multi-trait fine-mapping method flashfm [4], we also selected additional loci if they met the suggestive threshold p<10^-6^, with a second trait in the fine-mapping data set. As in Chen et al [1], the fine-mapping data-set did not include cohorts that were only genotyped with the Metabochip to avoid biasing results (**Supplementary table 1**).

Three loci *DPYSL5*, *GCKR* and *BRE* on chromosome 2 (coordinates chr2:26,640,022-27,677,428, chr2:27,230,940-28,242,603 and chr2:27,768,742-28,768,742) had overlapping coordinates suggesting shared causal variant(s). We merged them into a single locus *DPYSL5;GCKR;BRE* (coordinates chr2:26,730,940-29,268,742) and repeated single-ancestry fine-mapping.

### Reinserting key variants removed due to QC

Before performing multi-trait fine-mapping we first performed fine-mapping at 44 locus-trait associations where specific variants had previously been excluded from the analysis due to ancestry specific QC issues. As lead and index variants have a high probability of being causal due to high association evidence, we re-ran single-trait single-ancestry fine-mapping at 15 locus-trait associations at which index variants or top model variants were missing from any trait by including these variants back into the analysis for all traits associated with this locus (**Supplementary table 3**). Furthermore, multi-trait fine-mapping can only include variants present for all traits. To avoid the combination of variants (i.e. model) with the highest posterior probability (PP) of being causal being removed for some locus-trait associations, we also reinserted the top model variants from each of the traits associated with a locus into all of the other traits if not already present. Additionally, we repeated multi-ancestry fine-mapping at 29 locus-trait associations after re-inserting index and lead variants as well as EUR-like top model variants if the MPP was bigger than 0.8 in the same trait (**Supplementary table 3**). Four additional multi-ancestry locus-trait associations were missing EUR-like top model variants but were not rerun because either the likely causal gene at this locus was already identified (*PPARG/SYN2*, *TCF7L2, HFE*) or it was likely to affect HbA1c through non-glycaemic pathways (*SPTA1* and *HFE*).

### Single-trait fine-mapping with FINEMAP

After reinserting variants (see above), a shotgun stochastic search was carried out with FINEMAP v1.1 as in Chen et al [1]. Most parameters were kept as the default, except *--corr-config* was set to 0.9 and *--regions* to 1, to match the parameters used in Chen et al [1]. The maximum number of causal variants *n_c_* of each run was iteratively increased until the highest Bayes factor was achieved for *n_c_*-1 and the final version was run with n_c_-1 causal variants of the previous run i.e. the final version was run with the maximum number of causal variants set to the number of causal variants that had previously achieved the highest Bayes factor.

Chen et al [1] had used FINEMAP v1.1 for fine-mapping but only FINEMAP v1.4 allows for the addition of priors for each variant. To have a fair comparison of results, we reran the single-trait agnostic fine-mapping with FINEMAP v1.4 while keeping all the parameters the same. The default for parameter *--n-conv-sss* was 1000 in FINEMAP v1.1, but the new default in v1.4 is now set at 100. To keep all parameters the same, we changed this back to 1000. The maximum number of causal variants was set to the number determined by Chen et al [1] or to the maximum Bayes factor as determined with FINEMAP v1.1 for the rerun locus-trait associations.

Since multi-ancestry fine-mapping was carried out with FINEMAP v1.1 and compared to multi-trait fine-mapping using FINEMAP v1.4 single-trait results as input, we carried out a sensitivity analysis and compared multi-ancestry analysis (FINEMAP v1.1) with multi-trait analysis using the FINEMAP v1.1 single-trait analysis as input. This confirmed that the results were independent of the FINEMAP version used to generate the multi-trait fine-mapping results (**Supplementary Figure 5**).

### Multi-trait fine-mapping with flashfm

Flashfm (Hernández et al 2021) was used to perform multi-trait fine-mapping. Flashfm uses results from single-trait fine-mapping as input. Therefore, to conduct multi-trait fine-mapping single-trait fine-mapping results from FINEMAP v1.4 [1] were used.

Flashfm requires allele frequencies and effect sizes, which were taken from METAL results generated in Chen et al [1]. If FINEMAP and METAL defined different alleles as the reference allele, the frequency and effect sizes were adjusted to the FINEMAP reference allele. Up to four LD matrices were available for each locus, one for each associated trait. They were based on the subset of people that had the respective trait measured. Since not all trait measures were available for all individuals, the four matrices are based on different people and different sample sizes. For each locus, we used the LD matrix corresponding to the trait with the largest sample size that was also significantly associated with the locus. The LD matrix was converted to a covariance matrix using the reference allele frequency of the trait with the largest sample size. The flashfm parameters were left as the default apart from the odds of sharing between traits which was set to 80% (TOdds=0.25), to reflect the likely increased sharing of causal variants between related glycaemic traits. At the ADCY5 locus associated with all four traits, we noticed the CS size for HbA1c had increased from 8 in FINEMAP v1.1 to 2208 in FINEMAP v1.4. When running flashfm this led to the memory capacity being exceeded and fine-mapping failed at this locus with the default settings. Since there is at least one known causal variant (rs11708067) at the ADCY5 locus with functional support for involvement in insulin secretion [16], we excluded this locus for agnostic multi-trait fine-mapping.

### Functional enrichment analysis with fGWAS

For functional enrichment analysis, we considered the 60 annotations used by Chen et al [1] for all traits. We built a model of enriched annotations using fGWAS as described in Mahajan et al [17] and followed the suggested workflow of Pickrell [18]. For each trait we, started by determining the enrichment of all 60 annotations individually; annotations were defined as significantly enriched or depleted if the ln[parameter estimate] and corresponding 95% confidence interval were above or below zero, respectively. Of the significant annotations, we then selected the one with the highest natural logarithm of the maximum likelihood estimate of the enrichment parameter (llk) and added all remaining significant annotations in turn. Of these, we again selected the combination of annotations with the highest llk and kept adding all remaining significant annotations until none increased the llk. To avoid overfitting, we tested different cross-validation penalties in 0.05 increments and selected the one with the highest cross validation likelihood (cross-llk). We then dropped each of the annotations in the model in-turn while applying the cross-validation penalty and checked if any of the reduced models had an increased cross-llk. If so, the annotation was dropped and the process repeated until the highest cross-llk was reached and all estimates were significantly enriched according to their confidence interval. The remaining annotations and their estimates were then used to inform fine-mapping priors.

### Calculating variant priors for annotation-informed fine-mapping

We used the enrichment estimate of the parameters under the penalised likelihood (i.e. *β*) from the final model obtained from fGWAS as described above to calculate the priors for each SNP included in the fine-mapped loci. The prior for variant *i* is proportional to *Q_i_*

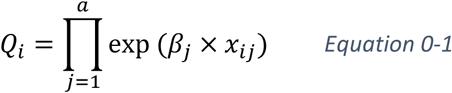

where *a* is the total number of annotations, *β_j_* is the enrichment estimate of annotation *j* and *x_ij_* is an indicator variable that takes 1 if variant *i* falls within annotation *j* and 0 if it does not.

The rescaled prior of variant *i* (*P_i_*) can then be calculated with

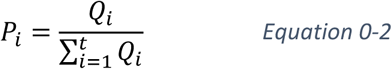

where *t* is the total number of variants at the locus.

### Single-trait annotation-informed fine-mapping

We carried out annotation-informed fine-mapping with FINEMAP v1.4 by providing it with the priors from fGWAS. The parameters and maximum number of causal variants were set as for the single-trait fine-mapping with FINEMAP v1.4.

### Multi-trait annotation-informed fine-mapping

For combined multi-trait annotation-informed fine-mapping we used the annotation-informed output from FINEMAP v1.4 as input for flashfm as detailed in Multi-trait fine-mapping with flashfm.

### Output parameters from fine-mapping results

For all fine-mapping results, we constructed 99% credible sets (99% CS) by summing up all models (containing one or more variants) from the largest to the smallest until they reached 99% posterior probability (PP) of containing or tagging the causal variants. The 99% CS size corresponds to the number of distinct variants in the 99% CS. Flashfm and FINEMAP calculate the marginal posterior probabilities of inclusion (MPP) of variants, which is the likelihood that a specific variant is causal or needed to tag the causal variant. This measure can take values between 0 and 1 with 1 indicating the variant has a 100% probability of being or tagging the causal variant.

### Data manipulation, visualisation and statistics

Data manipulation, visualisation and statistics were carried out in R [19]. The “tidyverse” [20] was used for data manipulation. The packages “ggpubr” [21] “ggsci” [22], “RColorBrewer” [23], “ggalluvial” [24], and “VennDiagram” [25] were used to generate figures. We used a two-sided paired Wilcoxon test from the package “rstatix” [26] to calculate p-values. P-values <0.05 were considered significant.

## Supporting information

Supplementary figures and MAGIC authors

## Data availability

Fine-mapping results (99% credible sets) will be available from the Downloads page of the Common Metabolic Disease Knowledge Portal (<u>cmdkp.org</u>) and the MAGIC website (https://www.magicinvestigators.org/).

## Code availability

Code will be available through Zenodo.

## Author contributions

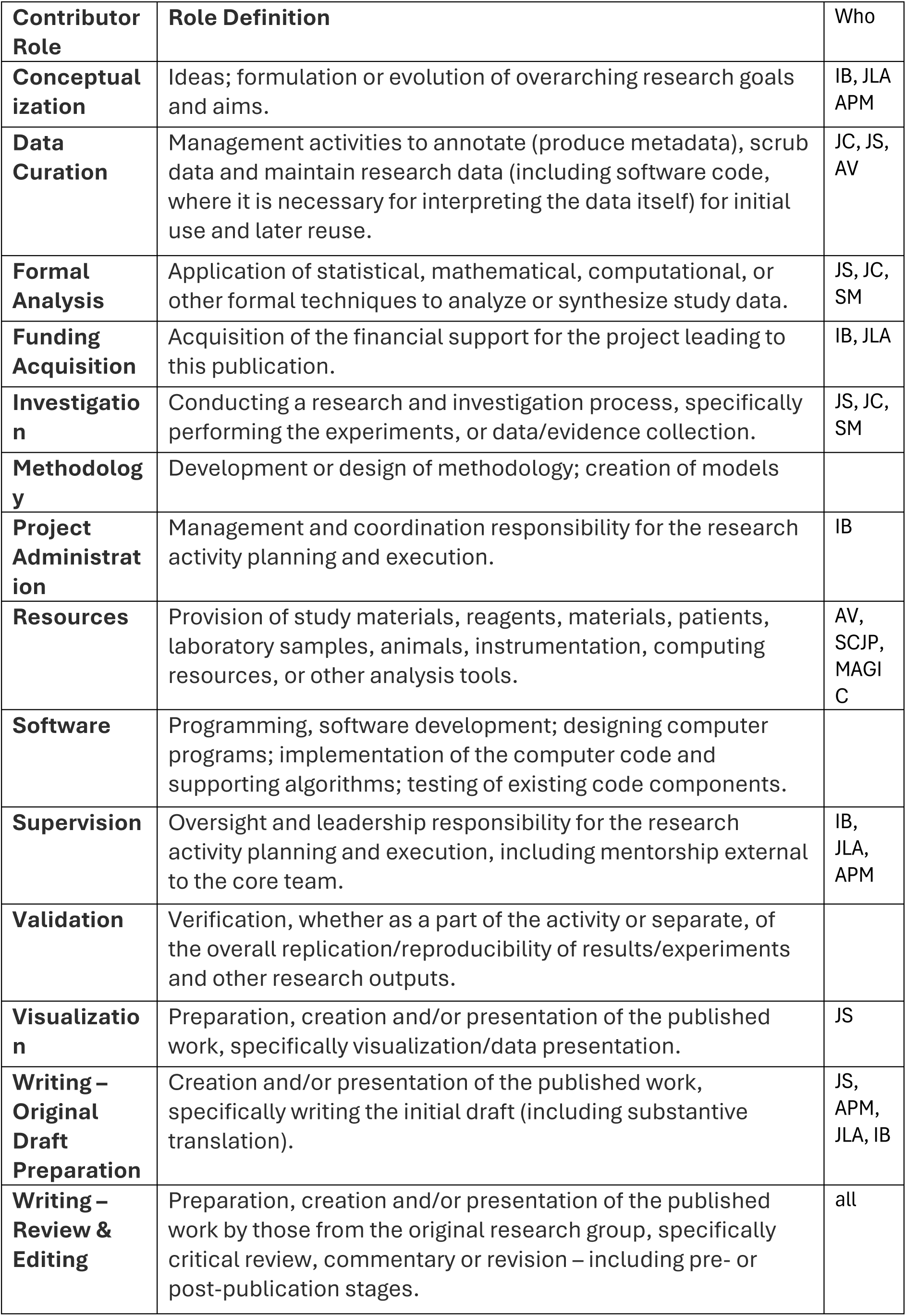

