## Supplementary figures and MAGIC authors for "Combining functional annotation and multi-trait fine-mapping methods improves fine-mapping resolution at glycaemic trait loci"

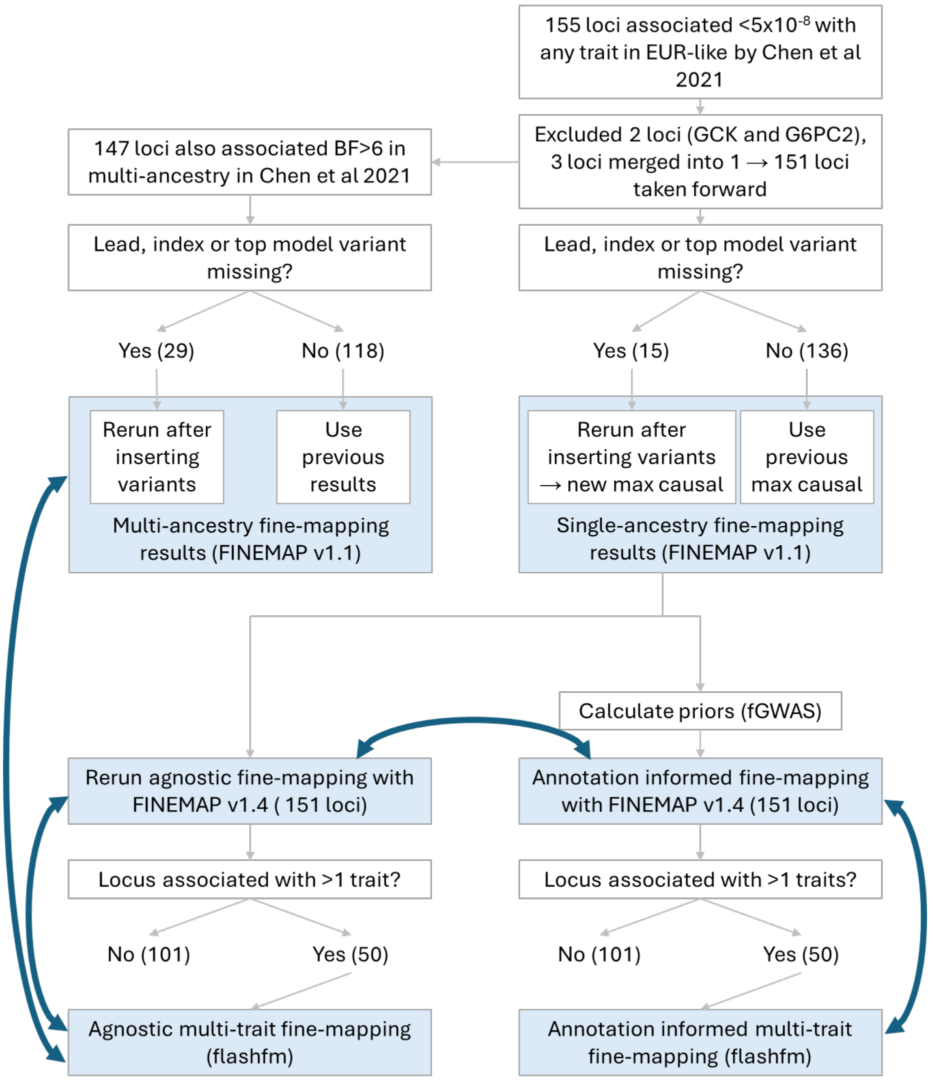


***Supplementary Figure 1 Study overview*** *We started with the fine-mapping results of 155 loci associated with glycaemic traits ($p<5x10^{-8}$) by Chen et al. We excluded two loci (GCK and G6PC2) which have multiple known causal variants per locus so fine-mapping would not resolve associations to a single variant and both genes have known roles in glucose metabolism. Another three overlapping loci were merged into a single locus. This left 151 loci that we considered for our study. Of these, 147 were also associated log10BF>6 in multi-ancestry analysis carried out with FINEMAP v1.1 by Chen et al. We re reinserted all lead and index variants from single and multi-ancestry analyses as well as top model variants from all traits associated with a locus. We then reran single-ancestry and multi-ancestry fine-mapping at the relevant loci. Chen et al had used FINEMAP v1.1 for fine-mapping but only v1.4 allows to incorporate priors from functional annotations. To allow for fair comparison, we re-ran 151 loci with FINEMAP v1.4 with the maximum number of causal variants determined by FINEMAP v1.1, either from previous results or after rerunning them with the reinserted variants. These were then used as the new agnostic single-trait fine-mapping results. Furthermore, there were 50 loci associated (p<10-6) with more than one trait which were fine-mapped with flashfm and their performance compared to single-trait fine-mapping as well as multi-ancestry fine-mapping. We also used fGWAS to build models of enriched annotations and calculate priors. These priors were used for annotation-informed fine-mapping with FINEMAP v1.4 and compared to agnostic fine-mapping with v1.4 We used these annotation-informed fine-mapping results as input for flashfm (at the 50 loci that were associated with more than one trait) for annotation-informed multi-trait fine-mapping. The results were compared to annotation-informed single-trait fine-mapping.*

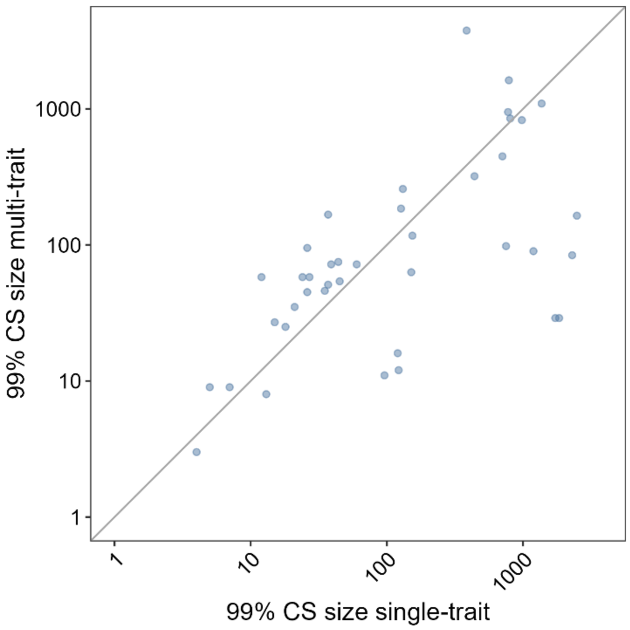


***Supplementary Figure 2 Multi-ancestry vs single-ancestry single-trait fine-mapping*** *Contains the subset of locus-trait associations where the multi-trait single-ancestry fine-mapping approach with flashfm performed better (smaller 99 % credible set (99% CS) size) than the single-trait multi-ancestry approach with FINEMAP v1.1. Each dot represents the 99% CS size at a locus-trait association according to single-trait fine-mapping with FINEMAP v1.4(x-axis), compared to single-trait multi-ancestry fine-mapping (y-axis) carried out with FINEMAP v1.1. The grey line represents the line of equality.*

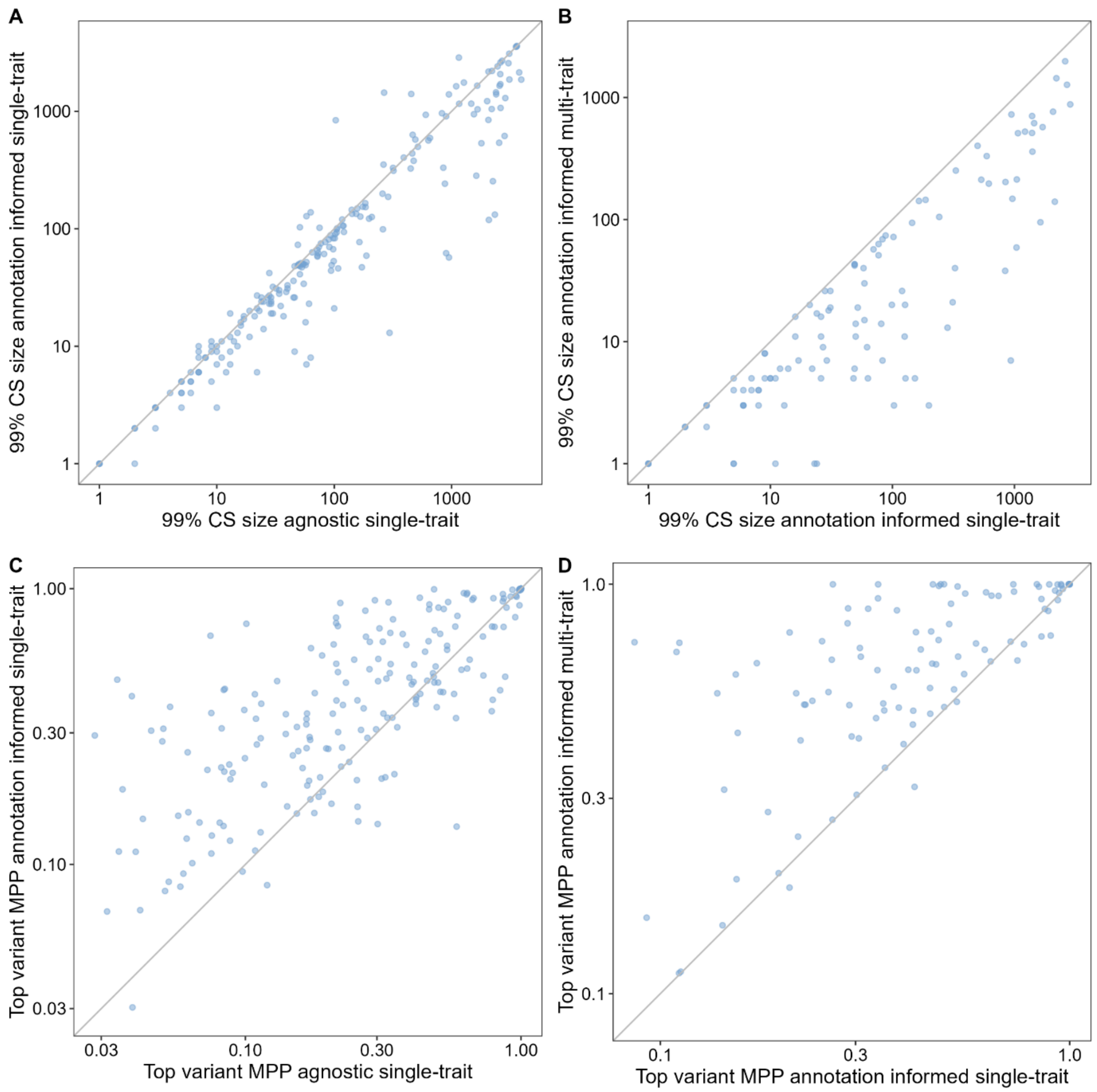


***Supplementary Figure 3 Comparison of agnostic and annotation-informed single-trait as well as multi-trait fine-mapping A&C****: Each dot represents the results at a locus-trait association according to agnostic fine-mapping (x-axis) compared to annotation-informed single-trait fine-mapping with FINEMAP v1.4 (y-axis) in EUR-like ancestry.* ***B&D*** *Compares annotation-informed single-trait fine-mapping with FINEMAP v1.4 (x-axis) to annotation-informed multi-trait fine-mapping (y-axis) with flashfm which uses the annotation-informed single-trait results as input. The grey line represents the line of equality. Prior probabilities for annotation-informed fine-mapping were obtained with fGWAS.* ***A&B****: Number of variants in the 99% credible set accounting for 99% of the posterior probability (PP) of variants being causal or tagging the causal variant.* ***C&D****: Marginal posterior probability (MPP) of variants with the highest MPP of each locus-trait association.*

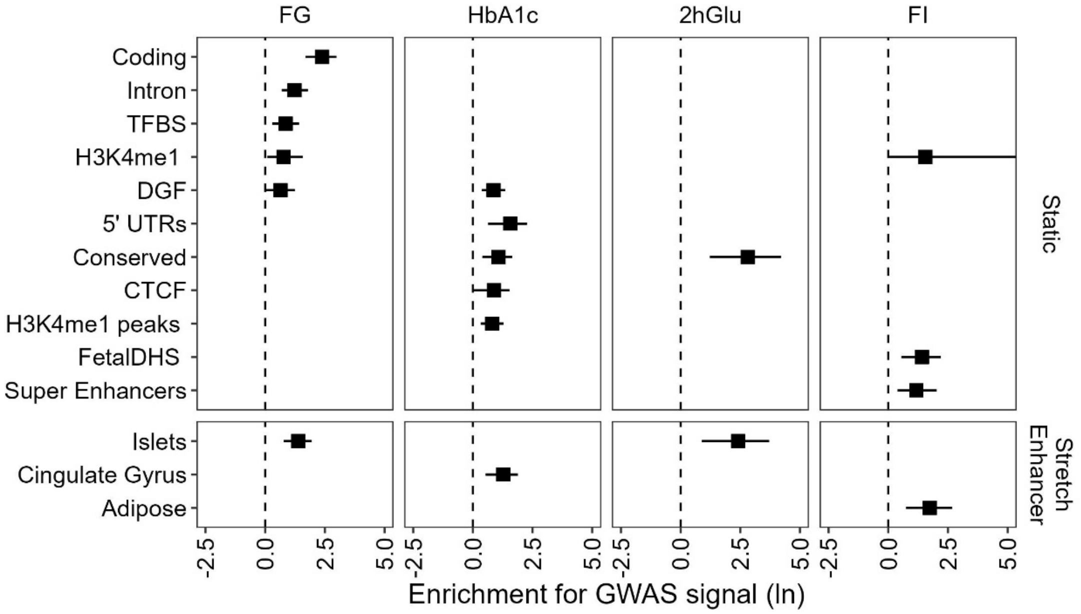


***Supplementary Figure 4 Joint model of enrichment of annotations in GWAS data.*** *Model best capturing the enrichment of annotations. Ln fold enrichment including 95% confidence interval from fGWAS. These estimates were used to calculate priors for annotation-informed fine-mapping.*

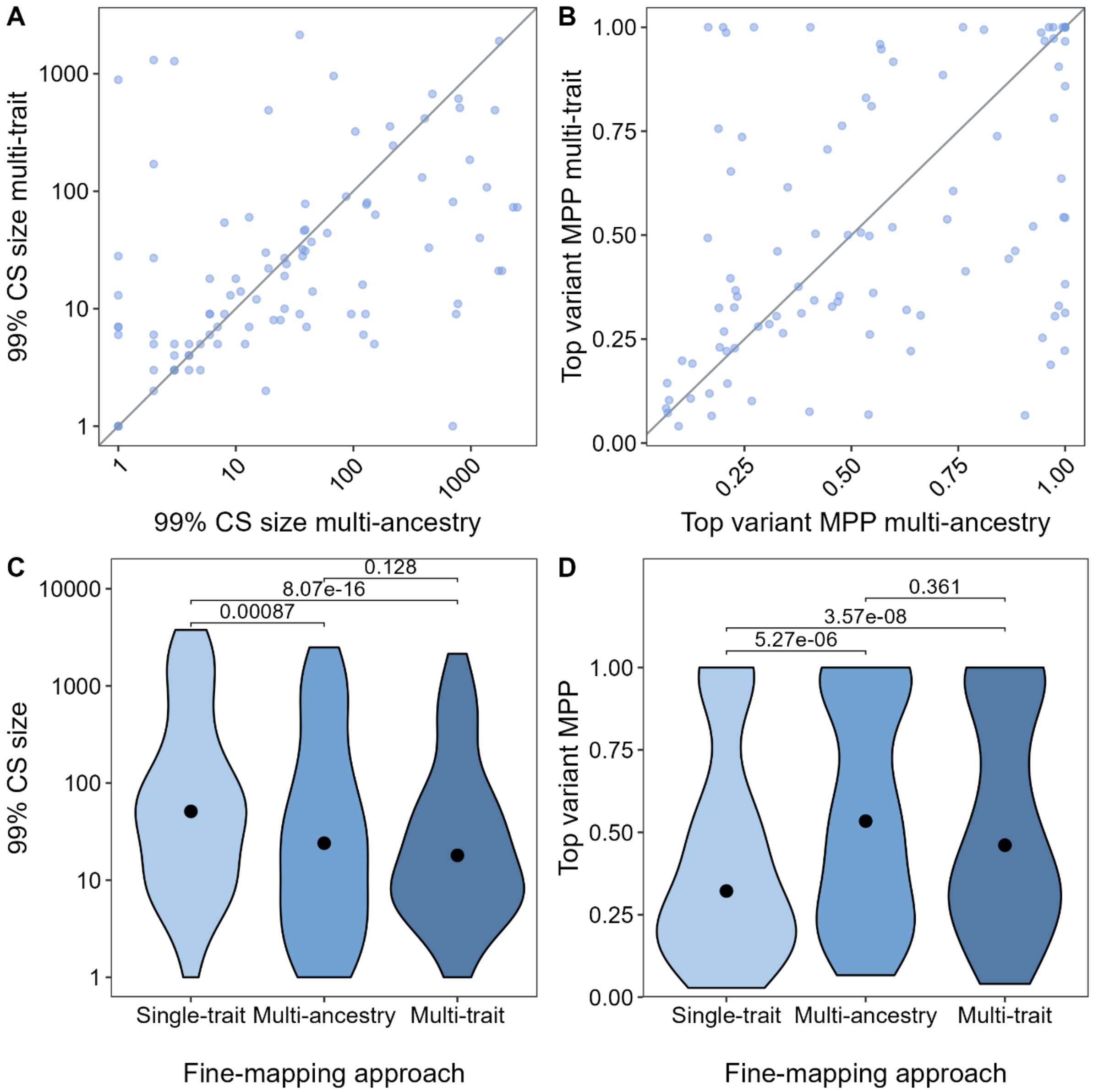


***Supplementary Figure 5 Comparison of agnostic fine-mapping approaches with FINEMAP v1.1*** *Multi-trait fine-mapping with flashfm in the main results section was carried out based on single-trait single-ancestry results from EUR-like data obtained by FINEMAP v1.4 which we compared to the original multi-ancestry single-trait fine-mapping with FINEMAP v1.1 from Chen et al. 2021. To check the version of FINEMAP did not influence these results we also compared multi-trait results based on FINEMAP v1.1 to multi-ancestry fine-mapping with FINEMAP v1.1.* ***A&B****: Each dot represents the results at a locus-trait association according to multi-ancestry (x-axis) compared to multi-trait approach (y-axis). The grey line represents the line of equality. The multi-trait results were obtained using the single-trait results from EUR-like as input.* ***C&D****: Includes only locus-trait associations that were analysed with all three approaches. The dot indicates the median, p-values were calculated with a paired two sided Wilcoxon.* ***A&C****: Compares the number of SNPs in the 99% credible set (99% CS).* ***B&D****: The marginal posterior probability (MPP) of the variant with the highest MPP of being causal.*

### MAGIC membership

Ji Chen[1,2,320], Cassandra N. Spracklen[3,4,320], Gaëlle Marenne [2,5,320], Arushi Varshney [6,320], Laura J. Corbin [7,8,320], Jian'an Luan[9], Sara M. Willems[9], Ying Wu[3], Xiaoshuai Zhang[9,10], Momoko Horikoshi[11,12,13], Thibaud S. Boutin[14], Reedik Mägi[15], Johannes Waage[16], Ruifang Li-Gao[17], Kei Hang Katie Chan[18,19,20], Jie Yao[21], Mila D. Anasanti[22], Audrey Y. Chu[23], Annique Claringbould[24], Jani Heikkinen[22], Jaeyoung Hong[25], Jouke-Jan Hottenga[26,27], Shaofeng Huo[28], Marika A. Kaakinen[22,29], Tin Louie[30], Winfried März[31,32,33], Hortensia Moreno-Macias[34], Anne Ndungu[12], Sarah C. Nelson[30], Ilja M. Nolte[35], Kari E. North[36], Chelsea K. Raulerson[3], Debashree Ray[37], Rebecca Rohde[36], Denis Rybin[25], Claudia Schurmann[38,39], Xueling Sim[40,41,42], Lorraine Southam[2,43], Isobel D. Stewart[9], Carol A. Wang[44], Yujie Wang[36], Peitao Wu[25], Weihua Zhang[45,46], Tarunveer S. Ahluwalia[16,47,48], Emil V. R. Appel[49], Lawrence F. Bielak[50], Jennifer A. Brody[51], Noël P. Burtt[52], Claudia P. Cabrera[53,54], Brian E. Cade[55,56], Jin Fang Chai[40], Xiaoran Chai[57,58], Li-Ching Chang[59], Chien-Hsiun Chen[59], Brian H. Chen[60], Kumaraswamy Naidu Chitrala[61], Yen-Feng Chiu[62], Hugoline G. de Haan[17], Graciela E. Delgado[33], Ayse Demirkan[29,63], Qing Duan[3,64], Jorgen Engmann[65], Segun A. Fatumo[66,67,68], Javier Gayán[69], Franco Giulianini[23], Jung Ho Gong[18], Stefan Gustafsson[70], Yang Hai[71], Fernando P. Hartwig[7,72], Jing He[73], Yoriko Heianza[74], Tao Huang[75], Alicia Huerta-Chagoya[76,77], Mi Yeong Hwang[78], Richard A. Jensen[51], Takahisa Kawaguchi[79], Katherine A. Kentistou[80,81], Young Jin Kim[78], Marcus E. Kleber[33], Ishminder K. Kooner[46], Shuiqing Lai[18], Leslie A. Lange[82], Carl D. Langefeld[83], Marie Lauzon[21], Man Li[84], Symen Ligthart[63], Jun Liu[63,85], Marie Loh[45,86], Jirong Long[87], Valeriya Lyssenko[88,89], Massimo Mangino[90,91], Carola Marzi[92,93], May E. Montasser[94], Abhishek Nag[12], Masahiro Nakatochi[95], Damia Noce[96], Raymond Noordam[97], Giorgio Pistis[98], Michael Preuss[38,99], Laura Raffield[3], Laura J. Rasmussen-Torvik[100], Stephen S. Rich[101,102], Neil R. Robertson[11,12], Rico Rueedi[103,104], Kathleen Ryan[94], Serena Sanna[24,98], Richa Saxena[105,106,107], Katharina E. Schraut[80,81], Bengt Sennblad[108], Kazuya Setoh[79], Albert V. Smith[109,110], Thomas Sparsø[49], Rona J. Strawbridge[111,112], Fumihiko Takeuchi[113], Jingyi Tan[21], Stella Trompet[97,114], Erik van den Akker[115,116,117], Peter J. van der Most[35], Niek Verweij[118,119], Mandy Vogel[120], Heming Wang[55,56], Chaolong Wang[121,122], Nan Wang[123,124], Helen R. Warren[53,54], Wanqing Wen[87], Tom Wilsgaard[125], Andrew Wong[126], Andrew R. Wood[1], Tian Xie[35], Mohammad Hadi Zafarmand[127,128], Jing-Hua Zhao[129], Wei Zhao[50], Najaf Amin[63,85], Zorayr Arzumanyan[21], Arne Astrup[130], Stephan J. L. Bakker[131], Damiano Baldassarre[132,133], Marian Beekman[115], Richard N. Bergman[134], Alain Bertoni[135], Matthias Blüher[136], Lori L. Bonnycastle[137], Stefan R. Bornstein[138], Donald W. Bowden[139], Qiuyin Cai[73], Archie Campbell[140,141], Harry Campbell[80], Yi Cheng Chang[59,142,143], Eco J. C. de Geus[26,27], Abbas Dehghan[63], Shufa Du[144], Gudny Eiriksdottir[110], Aliki Eleni Farmaki[145,146], Mattias Frånberg[112], Christian Fuchsberger[96], Yutang Gao[147], Anette P. Gjesing49, Anuj Goel12,148, Sohee Han78, Catharina A. Hartman149, Christian Herder[150,151,152], Andrew A. Hicks[96], Chang-Hsun Hsieh[153,154], Willa A. Hsueh[155], Sahoko Ichihara[156], Michiya Igase[157], M. Arfan Ikram[63], W. Craig Johnson[30], Marit E. Jørgensen[47,158], Peter K. Joshi[80], Rita R. Kalyani[159], Fouad R. Kandeel[160], Tomohiro Katsuya[161,162], Chiea Chuen Khor[122], Wieland Kiess[120], Ivana Kolcic[163], Teemu Kuulasmaa[164], Johanna Kuusisto[165], Kristi Läll[15], Kelvin Lam[21], Deborah A. Lawlor[7,8], Nanette R. Lee[166,167], Rozenn N. Lemaitre[51], Honglan Li[168], Lifelines Cohort Study*, Shih-Yi Lin[169,170], Jaana Lindström[171], Allan Linneberg[172,173], Jianjun Liu[122,174], Carlos Lorenzo[175], Tatsuaki Matsubara[176], Fumihiko Matsuda[79], Geltrude Mingrone[177], Simon Mooijaart[97], Sanghoon Moon[78], Toru Nabika[178], Girish N. Nadkarni[38], Jerry L. Nadler[179], Mari Nelis[15], Matt J. Neville[11,180], Jill M. Norris[181], Yasumasa Ohyagi[182], Annette Peters[93,183,184], Patricia A. Peyser[50], Ozren Polasek[163,185], Qibin Qi[186], Dennis Raven[149], Dermot F. Reilly[187], Alex Reiner[188], Fernando Rivideneira[189], Kathryn Roll[21], Igor Rudan[190], Charumathi Sabanayagam[57,191], Kevin Sandow[21], Naveed Sattar[192], Annette Schürmann[93,193], Jinxiu Shi[194], Heather M. Stringham[41,42], Kent D. Taylor[21], Tanya M. Teslovich[195], Betina Thuesen[172], Paul R. H. J.

Timmers[80,196], Elena Tremoli[133], Michael Y. Tsai[197], Andre Uitterlinden[189], Rob M. van Dam[40,174,198], Diana van Heemst[97], Astrid van Hylckama Vlieg[17], Jana V. van Vliet-Ostaptchouk[35], Jagadish Vangipurapu[199], Henrik Vestergaard[49,200], Tao Wang[186], Ko Willems van Dijk[201,202,203], Tatijana Zemunik[204], Gonçalo R. Abecasis[42], Linda S. Adair[144,205], Carlos Alberto Aguilar-Salinas[206,207,208], Marta E. Alarcón-Riquelme[209,210], Ping An[211], Larissa Aviles-Santa[212], Diane M. Becker[213], Lawrence J. Beilin[214], Sven Bergmann[103,104,215], Hans Bisgaard[16], Corri Black[216], Michael Boehnke[41,42], Eric Boerwinkle[217,218], Bernhard O. Böhm[219,220], Klaus Bønnelykke[16], D. I. Boomsma[26,27], Erwin P. Bottinger[38,221,222], Thomas A. Buchanan[124,223,224], Mickaël Canouil[225,226], Mark J. Caulfield[53,54], John C. Chambers[45,46,86,227,228], Daniel I. Chasman[23,229], Yii-Der Ida Chen[21], Ching-Yu Cheng[57,191], Francis S. Collins[137], Adolfo Correa[230], Francesco Cucca[98], H. Janaka de Silva[231], George Dedoussis[232], Sölve Elmståhl[233], Michele K. Evans[234], Ele Ferrannini[235], Luigi Ferrucci[236], Jose C. Florez[107,237,238], Paul W. Franks[89,239], Timothy M. Frayling[1], Philippe Froguel[225,226,240], Bruna Gigante[241], Mark O. Goodarzi[242], Penny Gordon-Larsen[144,205], Harald Grallert[92,93], Niels Grarup[49], Sameline Grimsgaard[125], Leif Groop[243,244], Vilmundur Gudnason[110,245], Xiuqing Guo[21], Anders Hamsten[112], Torben Hansen[49], Caroline Hayward[196], Susan R. Heckbert[246], Bernardo L. Horta[72], Wei Huang[194], Erik Ingelsson[247], Pankow S. James[248], Marjo-Ritta Jarvelin[249,250,251,252], Jost B. Jonas[253,254,255], J. Wouter Jukema[114,256], Pontiano Kaleebu[257], Robert Kaplan[186,188], Sharon L. R. Kardia[50], Norihiro Kato[113], Sirkka M. Keinanen-Kiukaanniemi[258,259], Bong-Jo Kim[78], Mika Kivimaki[260], Heikki A. Koistinen[261,262,263], Jaspal S. Kooner[46,227,228,264], Antje Körner[120], Peter Kovacs[136,265], Diana Kuh[126], Meena Kumari[266], Zoltan Kutalik[104,267], Markku Laakso[165], Timo A. Lakka[268,269,270], Lenore J. Launer[61], Karin Leander[271], Huaixing Li[28], Xu Lin[28], Lars Lind[272], Cecilia Lindgren[12,273,274], Simin Liu[18], Ruth J. F. Loos[38,99], Patrik K. E. Magnusson[275], Anubha Mahajan[12,319], Andres Metspalu[15], Dennis O. Mook-Kanamori[17,276], Trevor A. Mori[214], Patricia B. Munroe[53,54], Inger Njølstad[125], Jeffrey R. O'Connell[94], Albertine J. Oldehinkel[149], Ken K. Ong9, Sandosh Padmanabhan[277], Colin N. A. Palmer[278], Nicholette D. Palmer[139], Oluf Pedersen[49], Craig E. Pennell[44], David J. Porteous[140,279], Peter P. Pramstaller[96], Michael A. Province[211], Bruce M. Psaty[51,246,280], Lu Qi[281], Leslie J. Raffel[282], Rainer Rauramaa[270], Susan Redline[55,56], Paul M. Ridker[23,283], Frits R. Rosendaal[17], Timo E. Saaristo[284,285], Manjinder Sandhu[286], Jouko Saramies[287], Neil Schneiderman[288], Peter Schwarz[93,138,289], Laura J. Scott[41,42], Elizabeth Selvin[37], Peter Sever[264], Xiao-ou Shu[87], P. Eline Slagboom[115], Kerrin S. Small[90], Blair H. Smith[290], Harold Snieder[35], Tamar Sofer[238,291], Thorkild I. A. Sørensen[7,8,49,292], Tim D. Spector[90], Alice Stanton[293], Claire J. Steves[90,294], Michael Stumvoll[136], Liang Sun[28], Yasuharu Tabara[79], E. Shyong Tai[40,174,295], Nicholas J. Timpson[7,8], Anke Tönjes[136], Jaakko Tuomilehto[296,297,298], Teresa Tusie[77,299], Matti Uusitupa[300], Pim van der Harst[24,118], Cornelia van Duijn[63,85], Veronique Vitart[196], Peter Vollenweider[301], Tanja G. M. Vrijkotte[127], Lynne E. Wagenknecht[302], Mark Walker[303], Ya X. Wang[254], Nick J. Wareham9, Richard M. Watanabe[123,124,224], Hugh Watkins[12,148], Wen B. Wei[304], Ananda R. Wickremasinghe[305], Gonneke Willemsen[26,27], James F. Wilson[80,196], Tien-Yin Wong[57,191], Jer-Yuarn Wu[59], Anny H. Xiang[306], Lisa R. Yanek[213], Loïc Yengo[307], Mitsuhiro Yokota[308], Eleftheria Zeggini[2,43,309], Wei Zheng[87], Alan B. Zonderman[61], Jerome I. Rotter[21], Anna L. Gloyn[11,12,180,310], Mark I. McCarthy[11,12,180,311,319], Josée Dupuis[25], James B. Meigs[107,238,312], Robert A. Scott[9], Inga Prokopenko[22,29], Aaron Leong[229,313,314], Ching-Ti Liu[25], Stephen C. J. Parker[6,315,321], Karen L. Mohlke[3,321], Claudia Langenberg[9,321], Eleanor Wheeler[2,9,321], Andrew P. Morris[12,316,317,318,321], Inês Barroso[1,2,9,321]

### MAGIC affiliations

[1]Exeter Centre of Excellence for Diabetes Research (EXCEED), Genetics of Complex Traits, University of Exeter Medical School, University of Exeter, Exeter, UK. [2]Department of Human Genetics, Wellcome Sanger Institute, Cambridge, UK. [3]Department of Genetics, University of North Carolina, Chapel Hill, NC, USA. [4]Department of Biostatistics and Epidemiology, University of Massachusetts, Amherst, MA, USA. [5]Inserm, Univ Brest, EFS, UMR 1078, GGB, Brest, France. [6]Department of Computational Medicine and Bioinformatics, University of Michigan, Ann Arbor, MI, USA. [7]MRC Integrative Epidemiology Unit, University of Bristol, Bristol, UK. [8]Department of Population Health Sciences, Bristol Medical School, University of Bristol, Bristol, UK. [9]MRC Epidemiology Unit, Institute of Metabolic Science, University of Cambridge, Cambridge, UK. [10]Department of Biostatistics, School of Public Health, Shandong University, Jinan, China. [11]Oxford Centre for Diabetes, Endocrinology and Metabolism, Radcliffe Department of Medicine, University of Oxford, Oxford, UK. [12]Wellcome Centre for Human Genetics, University of Oxford, Oxford, UK. [13]Laboratory for Genomics of Diabetes and Metabolism, RIKEN Centre for Integrative Medical Sciences, Yokohama, Japan. [14]Medical Research Council Human Genetics Unit, Institute for Genetics and Molecular Medicine, Edinburgh, UK. [15]Estonian Genome Center, Institute of Genomics, University of Tartu, Tartu, Estonia. [16]COPSAC, Copenhagen Prospective Studies on Asthma in Childhood, Herlev and Gentofte Hospital, University of Copenhagen, Copenhagen, Denmark. [17]Department of Clinical Epidemiology, Leiden University Medical Center, Leiden, the Netherlands. [18]Department of Epidemiology, Brown University School of Public Health, Brown University, Providence, RI, USA. [19]Department of Biomedical Sciences, City University of Hong Kong, Hong Kong SAR, China. [20]Department of Electrical Engineering, City University of Hong Kong, Hong Kong SAR, China. [21]The Institute for Translational Genomics and Population Sciences, Department of Pediatrics, The Lundquist Institute for Biomedical Innovation at Harbor-UCLA Medical Center, Torrance, CA, USA. [22]Department of Metabolism, Digestion and Reproduction, Imperial College London, London, UK. [23]Division of Preventive Medicine, Brigham and Women's Hospital, Boston, MA, USA. [24]Department of Genetics, University of Groningen, University Medical Center Groningen, Groningen, the Netherlands. [25]Department of Biostatistics, Boston University School of Public Health, Boston, MA, USA. [26]Department of Biological Psychology, Faculty of Behaviour and Movement Sciences, Vrije Universiteit Amsterdam, Amsterdam, the Netherlands. [27]Amsterdam Public Health Research Institute, Amsterdam University Medical Center, Amsterdam, the Netherlands. [28]CAS Key Laboratory of Nutrition, Metabolism and Food Safety, Shanghai Institute of Nutrition and Health, University of Chinese Academy of Sciences, Chinese Academy of Sciences, Shanghai, China. [29]Section of Statistical Multi-omics, Department of Clinical and Experimental Research, University of Surrey, Guildford, UK. [30]Department of Biostatistics, University of Washington, Seattle, WA, USA. [31]SYNLAB Academy, SYNLAB Holding Deutschland GmbH, Mannheim, Germany. [32]Clinical Institute of Medical and Chemical Laboratory Diagnostics, Medical University Graz, Graz, Austria. [33]Vth Department of Medicine (Nephrology, Hypertensiology, Rheumatology, Endocrinology, Diabetology), Medical Faculty Mannheim, Heidelberg University, Mannheim, Baden-Württemberg, Germany. [34]Department of Economics, Metropolitan Autonomous University, Mexico City, Mexico. [35]Department of Epidemiology, University of Groningen, University Medical Center Groningen, Groningen, the Netherlands. [36]CVD Genetic Epidemiology Computational Laboratory, Gillings School of Global Public Health, University of North Carolina, Chapel Hill, NC, USA. [37]Department of Epidemiology, Johns Hopkins Bloomberg School of Public Health, Baltimore, MD, USA. [38]The Charles Bronfman Institute for Personalized Medicine, Icahn School of Medicine at Mount Sinai, New York, NY, USA. [39]HPI Digital Health

Center, Digital Health and Personalized Medicine, Hasso Plattner Institute, Potsdam, Germany. [40]Saw Swee Hock School of Public Health, National Univeristy of Singapore and National University Health System, Singapore, Singapore. [41]Center for Statistical Genetics, University of Michigan, Ann Arbor, MI, USA. [42]Department of Biostatistics, School of Public Health, University of Michigan, Ann Arbor, MI, USA. [43]Institute of Translational Genomics, Helmholtz Zentrum München–German Research Center for Environmental Health, Neuherberg, Germany. 44School of Medicine and Public Health, College of Health, Medicine and Wellbeing, The University of Newcastle, Newcastle, New South Wales, Australia. [45]Department of Epidemiology and Biostatistics, Imperial College London, London, UK. [46]Department of Cardiology, Ealing Hospital, London North West Healthcare NHS Trust, London, UK. [47]Steno Diabetes Center Copenhagen, Gentofte, Denmark. [48]The Bioinformatics Centre, Department of Biology, University of Copenhagen, Copenhagen, Denmark. [49]Novo Nordisk Foundation Center for Basic Metabolic Research, Faculty of Health and Medical Sciences, University of Copenhagen, Copenhagen, Denmark. [50]Department of Epidemiology, School of Public Health, University of Michigan, Ann Arbor, MI, USA. [51]Department of Medicine, Cardiovascular Health Research Unit, University of Washington, Seattle, WA, USA. [52]Metabolism Program, Program in Medical and Population Genetics, Broad Institute, Cambridge, MA, USA. [53]Department of Clinical Pharmacology, William Harvey Research Institute, Barts and The London School of Medicine and Dentistry, Queen Mary University of London, London, UK. [54]NIHR Barts Cardiovascular Biomedical Research Centre, Queen Mary University of London, London, UK. [55]Department of Medicine, Sleep and Circadian Disorders, Brigham and Women's Hospital, Boston, MA, USA. [56]Department of Medicine, Sleep Medicine, Harvard Medical School, Boston, MA, USA. [57]Ocular Epidemiology, Singapore Eye Research Institute, Singapore National Eye Centre, Singapore, Singapore. [58]Department of Ophthalmology, National University of Singapore and National University Health System, Singapore, Singapore. [59]Institute of Biomedical Sciences, Academia Sinica, Taipei, Taiwan. [60]Department of Epidemiology, The Herbert Wertheim School of Public Health and Human Longevity Science, University of California San Diego, La Jolla, CA, USA. [61]Laboratory of Epidemiology and Population Sciences, National Institute on Aging, National Institutes of Health, Baltimore, MD, USA. [62]Institute of Population Health Sciences, National Health Research Institutes, Miaoli, Taiwan. [63]Department of Epidemiology, Erasmus Medical Center, Rotterdam, the Netherlands. [64]Department of Statistics, University of North Carolina at Chapel Hill, Chapel Hill, NC, USA. [65]Institute of Cardiovascular Science, University College London, London, UK. [66]Uganda Medical Informatics Centre (UMIC), MRC/UVRI and London School of Hygiene & Tropical Medicine (Uganda Research Unit), Entebbe, Uganda. [67]London School of Hygiene & Tropical Medicine, London, UK. [68]H3Africa Bioinformatics Network (H3ABioNet) Node, Centre for Genomics Research and Innovation, NABDA/FMST, Abuja, Nigeria. [69]Bioinfosol, Sevilla, Spain. [70]Molecular Epidemiology and Science for Life Laboratory, Department of Medical Sciences, Uppsala University, Uppsala, Sweden. [71]Department of Statistics, The University of Auckland, Science Center, Auckland, New Zealand. [72]Postgraduate Program in Epidemiology, Federal University of Pelotas, Pelotas, Brazil. [73]Department of Medicine, Epidemiology, Vanderbilt University Medical Center, Nashville, TN, USA. [74]Department of Epidemiology, Tulane University Obesity Research Center, Tulane University, New Orleans, LA, USA. [75]Department of Epidemiology and Biostatistics, School of Public Health, Peking University, Beijing, China. [76]Molecular Biology and Genomic Medicine Unit, National Council for Science and Technology, Mexico City, Mexico. [77]Molecular Biology and Genomic Medicine Unit, National Institute of Medical Sciences and Nutrition, Mexico City, Mexico. [78]Division of Genome Science, Department of Precision Medicine, National Institute of Health, Cheongju, South Korea. [79]Center for Genomic Medicine, Kyoto University Graduate School of Medicine, Kyoto, Japan. [80]Centre for Global

Health Research, Usher Institute, University of Edinburgh, Edinburgh, UK. [81]Centre for Cardiovascular Sciences, Queen's Medical Research Institute, University of Edinburgh, Edinburgh, UK. [82]Department of Medicine, Divison of Biomedical Informatics and Personalized Medicine, University of Colorado Anschutz Medical Campus, Denver, CO, USA. [83]Department of Biostatistics and Data Science, Wake Forest School of Medicine, Winston-Salem, NC, USA. [84]Department of Medicine, Division of Nephrology and Hypertension, University of Utah, Salt Lake City, UT, USA. [85]Nuffield Department of Population Health, University of Oxford, Oxford, UK. [86]Lee Kong Chian School of Medicine, Nanyang Technological University, Singapore, Singapore. [87]Division of Epidemiology, Department of Medicine, Vanderbilt Epidemiology Center, Vanderbilt University Medical Center, Nashville, TN, USA. [88]Department of Clinical Science, Center for Diabetes Research, University of Bergen, Bergen, Norway. [89]Department of Clinical Sciences, Lund University Diabetes Centre, Lund University, Malmo, Sweden. [90]Department of Twin Research and Genetic Epidemiology, School of Life Course Sciences, King's College London, London, UK. [91]NIHR Biomedical Research Centre, Guy's and St Thomas' NHS Foundation Trust, London, UK. [92]Institute of Epidemiology, Research Unit of Molecular Epidemiology, Helmholtz Zentrum München Research Center for Environmental Health, Neuherberg, Germany. [93]German Center for Diabetes Research (DZD), Neuherberg, Germany. [94]Department of Medicine, Division of Endocrinology, Diabetes and Nutrition, University of Maryland School of Medicine, Baltimore, MD, USA. [95]Public Health Informatics Unit, Department of Integrated Sciences, Nagoya University Graduate School of Medicine, Nagoya, Japan. [96]Institute for Biomedicine, Eurac Research, Bolzano, Italy. [97]Department of Internal Medicine, Section of Gerontology and Geriatrics, Leiden University Medical Center, Leiden, the Netherlands. [98]Istituto di Ricerca Genetica e Biomedica (IRGB), Consiglio Nazionale delle Ricerche (CNR), Monserrato, Italy. [99]The Mindich Child Health and Development Institute for Personalized Medicine, Icahn School of Medicine at Mount Sinai, New York, NY, USA. [100]Department of Preventive Medicine, Northwestern University Feinberg School of Medicine, Chicago, IL, USA. [101]Center for Public Health Genomics, University of Virginia, Charlottesville, VA, USA. [102]Department of Public Health Sciences, University of Virginia, Charlottesville, VA, USA. [103]Department of Computational Biology, University of Lausanne, Lausanne, Switzerland. [104]Swiss Institute of Bioinformatics, Lausanne, Switzerland. [105]Center for Genomic Medicine, Massachusetts General Hospital, Harvard Medical School, Boston, MA, USA. [106]Department of Anesthesia, Critical Care and Pain Medicine, Massachusetts General Hospital, Boston, MA, USA. [107]Program in Medical and Population Genetics, Broad Institute, Cambridge, MA, USA. [108]Department of Cell and Molecular Biology, National Bioinformatics Infrastructure Sweden, Science for Life Laboratory, Uppsala University, Uppsala, Sweden. [109]Department of Biostatistics, University of Michigan, Ann Arbor, MI, USA. [110]Icelandic Heart Association, Kopavogur, Iceland. [111]Institute of Health and Wellbeing, University of Glasgow, Glasgow, UK. [112]Department of Medicine Solna, Cardiovascular Medicine, Karolinska Institutet, Stockholm, Sweden. [113]National Center for Global Health and Medicine, Tokyo, Japan. [114]Department of Cardiology, Leiden University Medical Center, Leiden, the Netherlands. [115]Department of Biomedical Data Sciences, Molecular Epidemiology, Leiden University Medical Center, Leiden, the Netherlands. [116]Department of Pattern Recognition and Bioinformatics, Delft University of Technology, Delft, the Netherlands. [117]Department of Biomedical Data Sciences, Leiden Computational Biology Center, Leiden University Medical Center, Leiden, the Netherlands. [118]Department of Cardiology, University of Groningen, University Medical Center Groningen, Groningen, the Netherlands. [119]Genomics PLC, Oxford, UK. [120]Center of Pediatric Research, University Children's Hospital Leipzig, University of Leipzig Medical Center, Leipzig, Germany. [121]Department of Epidemiology and Biostatistics, School of Public Health, Tongji Medical College, Huazhong University of Science and Technology, Wuhan,

China. [122]Genome Institute of Singapore, Agency for Science, Technology and Research, Singapore, Singapore. [123]Department of Preventive Medicine, Keck School of Medicine of University of Southern California, Los Angeles, CA, USA. [124]University of Southern California Diabetes and Obesity Research Institute, Keck School of Medicine of University of Southern California, Los Angeles, CA, USA. [125]Department of Community Medicine, Faculty of Health Sciences, UIT the Arctic University of Norway, Tromsø, Norway. [126]MRC Unit for Lifelong Health and Ageing at University College London, London, UK. [127]Department of Public Health, Amsterdam Public Health Research Institute, Amsterdam University Medical Center, Amsterdam, the Netherlands. [128]Department of Clinical Epidemiology, Biostatistics, and Bioinformatics, Amsterdam Public Health Research Institute, Amsterdam University Medical Center, Amsterdam, the Netherlands. [129]Department of Public Health and Primary Care, School of Clinical Medicine, University of Cambridge, Cambridge, UK. [130]Department of Nutrition, Exercise, and Sports, Faculty of Science, University of Copenhagen, Copenhagen, Denmark. [131]Department of Internal Medicine, University of Groningen, University Medical Center Groningen, Groningen, the Netherlands. [132]Department of Medical Biotechnology and Translational Medicine, University of Milan, Milan, Italy. [133]Centro Cardiologico Monzino, IRCCS, Milan, Italy. [134]Diabetes and Obesity Research Institute, Cedars-Sinai Medical Center, Los Angeles, CA, USA. [135]Department of Epidemiology and Prevention, Division of Public Health Sciences, Wake Forest School of Medicine, Winston-Salem, NC, USA. [136]Medical Department III–Endocrinology, Nephrology, Rheumatology, University of Leipzig Medical Center, Leipzig, Germany. [137]Medical Genomics and Metabolic Genetics Branch, National Human Genome Research Institute, National Institues of Health, Bethesda, MD, USA. [138]Department for Prevention and Care of Diabetes, Faculty of Medicine Carl Gustav Carus, Technische Universität Dresden, Dresden, Germany. [139]Department of Biochemistry, Wake Forest School of Medicine, Winston-Salem, NC, USA. [140]Centre for Genomic and Experimental Medicine, Institute of Genetics and Molecular Medicine, University of Edinburgh, Western General Hospital, Edinburgh, UK. [141]Usher Institute, University of Edinburgh, Edinburgh, UK. [142]Department of Internal Medicine, National Taiwan University Hospital, Taipei, Taiwan. [143]Graduate Institute of Medical Genomics and Proteomics, National Taiwan University, Taipei, Taiwan. [144]Department of Nutrition, Gillings School of Global Public Health, University of North Carolina, Chapel Hill, NC, USA. [145]Department of Population Science and Experimental Medicine, Institute of Cardiovascular Science, University College London, London, UK. [146]Department of Nutrition and Dietetics, School of Health Science and Education, Harokopio University of Athens, Athens, Greece. [147]Department of Epidemiology, Shanghai Cancer Institute, Shanghai, China. [148]Division of Cardiovascular Medicine, Radcliffe Department of Medicine, University of Oxford, Oxford, UK. [149]Department of Psychiatry, Interdisciplinary Center Psychopathy and Emotion Regulation, University of Groningen, University Medical Center Groningen, Groningen, the Netherlands. [150]Institute for Clinical Diabetology, German Diabetes Center, Leibniz Center for Diabetes Research at Heinrich Heine University Düsseldorf, Düsseldorf, Germany. [151]Division of Endocrinology and Diabetology, Medical Faculty, Heinrich Heine University Düsseldorf, Düsseldorf, Germany. [152]German Center for Diabetes Research (DZD), Düsseldorf, Germany. [153]Internal Medicine, Endocrine and Metabolism, Tri-Service General Hospital, Taipei, Taiwan. [154]School of Medicine, National Defense Medical Center, Taipei, Taiwan. [155]Internal Medicine, Endocrinology, Diabetes and Metabolism, Diabetes and Metabolism Research Center, The Ohio State University Wexner Medical Center, Columbus, OH, USA. [156]Department of Environmental and Preventive Medicine, Jichi Medical University School of Medicine, Shimotsuke, Japan. [157]Department of Anti-aging Medicine, Ehime University Graduate School of Medicine, Toon, Japan. [158]National Institute of Public Health, University of Southern Denmark, Odense, Denmark.

[159]Department of Medicine, Endocrinology, Diabetes and Metabolism, Johns Hopkins University School of Medicine, Baltimore, MD, USA. [160]Clinical Diabetes, Endocrinology and Metabolism, Translational Research and Cellular Therapeutics, Beckman Research Institute of the City of Hope, Duarte, CA, USA. [161]Department of Clinical Gene Therapy, Osaka University Graduate School of Medicine, Suita, Japan. [162]Department of Geriatric and General Medicine, Osaka University Graduate School of Medicine, Suita, Japan. [163]Department of Public Health, University of Split School of Medicine, Split, Croatia. [164]Institute of Biomedicine, Bioinformatics Center, Univeristy of Eastern Finland, Kuopio, Finland. [165]Department of Medicine, University of Eastern Finland and Kuopio University Hospital, Kuopio, Finland. [166]USC-Office of Population Studies Foundation, University of San Carlos, Cebu City, the Philippines. [167]Department of Anthropology, Sociology and History, University of San Carlos, Cebu City, the Philippines. [168]State Key Laboratory of Oncogene and Related Genes and Department of Epidemiology, Shanghai Cancer Institute, Renji Hospital, Shanghai Jiaotong University School of Medicine, Shanghai, China. [169]Center for Geriatrics and Gerontology, Taichung Veterans General Hospital, Taichung, Taiwan. [170]National Defense Medical Center, National Yang-Ming University, Taipei, Taiwan. [171]Diabetes Prevention Unit, National Institute for Health and Welfare, Helsinki, Finland. [172]Center for Clinical Research and Prevention, Bispebjerg and Frederiksberg Hospital, Copenhagen, Denmark. [173]Department of Clinical Medicine, Faculty of Health and Medical Sciences, University of Copenhagen, Copenhagen, Denmark. [174]Yong Loo Lin School of Medicine, National University of Singapore and National University Health System, Singapore, Singapore. [175]Department of Medicine, University of Texas Health Sciences Center, San Antonio, TX, USA. [176]Department of Internal Medicine, Aichi Gakuin University School of Dentistry, Nagoya, Japan. [177]Department of Diabetes, Diabetes, and Nutritional Sciences, James Black Centre, King's College London, London, UK. [178]Department of Functional Pathology, Shimane University School of Medicine, Izumo, Japan. [179]Department of Medicine and Pharmacology, New York Medical College School of Medicine, Valhalla, NY, USA. [180]Oxford NIHR Biomedical Research Centre, Oxford University Hospitals NHS Foundation Trust, Oxford, UK. [181]Colorado School of Public Health, University of Colorado Anschutz Medical Campus, Aurora, CO, USA. [182]Department of Geriatric Medicine and Neurology, Ehime University Graduate School of Medicine, Toon, Japan. [183]Institute of Epidemiology, Helmholtz Zentrum München Research Center for Environmental Health, Neuherberg, Germany. [184]Institute for Medical Information Processing, Biometry and Epidemiology, Ludwig-Maximilians University Munich, Munich, Germany. [185]Gen-Info, Zagreb, Croatia. [186]Department of Epidemiology and Population Health, Albert Einstein College of Medicine, New York, NY, USA. [187]Genetics and Pharmacogenomics, Merck Sharp & Dohme, Kenilworth, NJ, USA. [188]Department of Public Health Sciences, Fred Hutchinson Cancer Research Center, Seattle, WA, USA. [189]Department of Internal Medicine, Erasmus Medical Center, Rotterdam, the Netherlands. [190]Centre for Global Health, The Usher Institute, University of Edinburgh, Edinburgh, UK. [191]Ophthalmology & Visual Sciences Academic Clinical Program (Eye ACP), Duke-NUS Medical School, Singapore, Singapore. [192]BHF Glasgow Cardiovascular Research Centre, Institute of Cardiovascular and Medical Sciences, University of Glasgow, Glasgow, UK. [193]Department of Experimental Diabetology, German Institute of Human Nutrition Potsdam-Rehbruecke, Nuthetal, Germany. [194]Department of Genetics, Shanghai-MOST Key Laboratory of Health and Disease Genomics, Chinese National Human Genome Center at Shanghai (CHGC) and Shanghai Academy of Science & Technology (SAST), Shanghai, China. [195]Sarepta Therapeutics, Cambridge, MA, USA. [196]Medical Research Council Human Genetics Unit, Institute for Genetics and Cancer, University of Edinburgh, Edinburgh, UK. [197]Department of Laboratory Medicine and Pathology, University of Minnesota, Minneapolis, MN, USA. [198]Department of Nutrition, Harvard T. H. Chan School of Public Health, Boston, MA, USA.

[199]Institute of Clinical Medicine, Internal Medicine, University of Eastern Finland, Kuopio, Finland. [200]Department of Medicine, Bornholms Hospital, Rønne, Denmark. [201]Department of Internal Medicine, Division of Endocrinology, Leiden University Medical Center, Leiden, the Netherlands. [202]Laboratory for Experimental Vascular Medicine, Leiden University Medical Center, Leiden, the Netherlands. [203]Department of Human Genetics, Leiden University Medical Center, Leiden, the Netherlands. [204]Department of Human Biology, University of Split School of Medicine, Split, Croatia. [205]Carolina Population Center, University of North Carolina, Chapel Hill, NC, USA. [206]Department of Endocrinology and Metabolism, Instituto Nacional de Ciencias Medicas y Nutricion, Mexico City, Mexico. [207]Unidad de Investigación de Enfermedades Metabólicas, Instituto Nacional de Ciencias Médicas y Nutrición and Tec Salud, Mexico City, Mexico. [208]Instituto Tecnológico y de Estudios Superiores de Monterrey Tec Salud, Monterrey, Mexico. [209]Department of Medical Genomics, Pfizer/University of Granada/Andalusian Government Center for Genomics and Oncological Research (GENYO), Granada, Spain. [210]Institute for Environmental Medicine, Chronic Inflammatory Diseases, Karolinska Institutet, Solna, Sweden. [211]Department of Genetics, Division of Statistical Genomics, Washington University School of Medicine, St Louis, MO, USA. [212]Clinical and Health Services Research, National Institute on Minority Health and Health Disparities, Bethesda, MD, USA. [213]Department of Medicine, General Internal Medicine, Johns Hopkins University School of Medicine, Baltimore, MD, USA. [214]Medical School, Royal Perth Hospital Unit, University of Western Australia, Perth, Western Australia, Australia. [215]Department of Integrative Biomedical Sciences, University of Cape Town, Cape Town, South Africa. [216]Aberdeen Centre for Health Data Science, School of Medicine, Medical Sciences and Nutrition, University of Aberdeen, Aberdeen, UK. [217]Human Genetics Center, School of Public Health, The University of Texas Health Science Center at Houston, Houston, TX, USA. [218]Human Genome Sequencing Center, Baylor College of Medicine, Houston, TX, USA. [219]Division of Endocrinology and Diabetes, Graduate School of Molecular Endocrinology and Diabetes, University of Ulm, Ulm, Germany. [220]LKC School of Medicine, Nanyang Technological University, Singapore and Imperial College London, UK, Singapore, Singapore. [221]Hasso Plattner Institute for Digital Health at Mount Sinai, Icahn School of Medicine at Mount Sinai, New York, NY, USA. [222]Digital Health Center, Hasso Plattner Institut, University Potsdam, Potsdam, Germany. [223]Department of Medicine, Keck School of Medicine of University of Southern California, Los Angeles, CA, USA. [224]Department of Physiology and Neuroscience, Keck School of Medicine of University of Southern California, Los Angeles, CA, USA. [225]INSERM UMR 1283/CNRS UMR 8199, European Institute for Diabetes (EGID), Université de Lille, Lille, France. [226]INSERM UMR 1283/CNRS UMR 8199, European Institute for Diabetes (EGID), Institut Pasteur de Lille, Lille, France. [227]Imperial College Healthcare NHS Trust, Imperial College London, London, UK. [228]MRC-PHE Centre for Environment and Health, Imperial College London, London, UK. [229]Harvard Medical School, Boston, MA, USA. [230]Department of Medicine, Jackson Heart Study, University of Mississippi Medical Center, Jackson, MS, USA. [231]Department of Medicine, Faculty of Medicine, University of Kelaniya, Ragama, Sri Lanka. [232]Department of Nutrition and Dietetics, School of Health Science and Education, Harokopio University of Athens, Kallithea, Greece. [233]Department of Clinical Sciences, Lund University, Malmö, Sweden. [234]Laboratory of Epidemiology and Population Sciences, National Institute on Aging Intramural Research Program, National Institutes of Health, Baltimore, MD, USA. [235]CNR Institute of Clinical Physiology, Pisa, Italy. [236]Intramural Research Program, National Institute of Aging, Baltimore, MD, USA. [237]Diabetes Unit and Center for Genomic Medicine, Massachusetts General Hospital, Boston, MA, USA. [238]Department of Medicine, Harvard Medical School, Boston, MA, USA. [239]Department of Public Health and Clinical Medicine, Umeå University, Umeå, Sweden. [240]Department of Genomics of Common Disease, Imperial College London, London, UK.

[241]Department of Medicine, Cardiovascular Medicine, Karolinska Institutet, Stockholm, Sweden. [242]Department of Medicine, Division of Endocrinology, Diabetes and Metabolism, Cedars-Sinai Medical Center, Los Angeles, CA, USA. [243]Diabetes Centre, Lund University, Lund, Sweden. [244]Finnish Institute of Molecular Medicine, Helsinki University, Helsinki, Finland. [245]Faculty of Medicine, School of Health Sciences, University of Iceland, Reykjavik, Iceland. [246]Department of Epidemiology, Cardiovascular Health Research Unit, University of Washington, Seattle, WA, USA. [247]Department of Medicine, Division of Cardiovascular Medicine, Stanford University School of Medicine, Stanford University, Stanford, CA, USA. [248]Division of Epidemiology and Community Health, University of Minnesota, Minneapolis, MN, USA. [249]Department of Epidemiology and Biostatistics, MRC-PHE Centre for Environment and Health, School of Public Health, Imperial College London, London, UK. [250]Center for Life Course Health Research, Faculty of Medicine, University of Oulu, Oulu, Finland. [251]Unit of Primary Health Care, Oulu Univerisity Hospital, OYS, Oulu, Finland. [252]Department of Life Sciences, College of Health and Life Sciences, Brunel University London, London, UK. [253]Department of Ophthalmology, Medical Faculty Mannheim, Heidelberg University, Mannheim, Germany. [254]Beijing Institute of Ophthalmology, Beijing Ophthalmology and Visual Science Key Lab, Beijing Tongren Eye Center, Beijing Tongren Hospital, Capital Medical University, Beijing, China. [255]Institute of Molecular and Clinical Ophthalmology Basel IOB, Basel, Switzerland. [256]Netherlands Heart Institute, Utrecht, the Netherlands. [257]MRC/UVRI and LSHTM (Uganda Research Unit), Entebbe, Uganda. [258]Faculty of Medicine, Institute of Health Sciences, University of Oulu, Oulu, Finland. [259]Unit of General Practice, Oulu University Hospital, Oulu, Finland. [260]Department of Epidemiology and Public Health, University College London, London, UK. [261]Department of Public Health Solutions, Finnish Institute for Health and Welfare, Helsinki, Finland. [262]Department of Medicine, University of Helsinki and Helsinki University Central Hospital, Helsinki, Finland. [263]Minerva Foundation Institute for Medical Research, Helsinki, Finland. [264]National Heart and Lung Institute, Imperial College London, London, UK. [265]IFB Adiposity Diseases, University of Leipzig Medical Center, Leipzig, Germany. [266]Institute for Social and Economic Research, University of Essex, Colchester, UK. [267]University Institute of Primary Care and Public Health, Division of Biostatistics, University of Lausanne, Lausanne, Switzerland. [268]Institute of Biomedicine, School of Medicine, University of Eastern Finland, Kuopio, Finland. [269]Department of Clinical Physiology and Nuclear Medicine, Kuopio University Hospital, Kuopio, Finland. [270]Foundation for Research in Health Exercise and Nutrition, Kuopio Research Institute of Exercise Medicine, Kuopio, Finland. [271]Institute of Environmental Medicine, Cardiovascular and Nutritional Epidemiology, Karolinska Institutet, Stockholm, Sweden. [272]Department of Medical Sciences, University of Uppsala, Uppsala, Sweden. [273]Big Data Institute, Nuffield Department of Medicine, University of Oxford, Oxford, UK. [274]Nuffield Department of Women's and Reproductive Health, University of Oxford, Oxford, UK. [275]Department of Medical Epidemiology and Biostatistics and the Swedish Twin Registry, Karolinska Institutet, Stockholm, Sweden. [276]Department of Public Health and Primary Care, Leiden University Medical Center, Leiden, the Netherlands. [277]Institute of Cardiovascular and Medical Sciences, University of Glasgow, Glasgow, UK. [278]Division of Population Health and Genomics, School of Medicine, University of Dundee, Ninewells Hospital and Medical School, Dundee, UK. [279]Centre for Cognitive Ageing and Cognitive Epidemiology, University of Edinburgh, Edinburgh, UK. [280]Department of Health Services, Cardiovascular Health Research Unit, University of Washington, Seattle, WA, USA. [281]Department of Epidemiology, Tulane University School of Public Health and Tropical Medicine, New Orleans, LA, USA. [282]Department of Pediatrics, Genetic and Genomic Medicine, University of California, Irvine, Irvine, CA, USA. [283]Havard Medical School, Boston, MA, USA. [284]Tampere, Finnish Diabetes Association, Tampere, Finland. [285]Pirkanmaa Hospital District, Tampere, Finland. [286]Department of Medicine, University

of Cambridge, Cambridge, UK. [287]South Karelia Central Hospital, Lappeenranta, Finland. [288]Department of Psychology, University of Miami, Miami, FL, USA. [289]Paul Langerhans Institute Dresden of the Helmholtz Center Munich, University Hospital and Faculty of Medicine, Dresden, Germany. [290]Division of Population Health and Genomics, Ninewells Hospital and Medical School, University of Dundee, Dundee, UK. [291]Division of Sleep and Circadian Disorders, Brigham and Women's Hospital, Boston, MA, USA. [292]Department of Public Health, Section of Epidemiology, Faculty of Health and Medical Sciences, University of Copenhagen, Copenhagen, Denmark. [293]Department of Molecular and Cellular Therapeutics, Royal College of Surgeons in Ireland, Dublin, Ireland. [294]Department of Ageing and Health, Guy's and St Thomas' NHS Foundation Trust, London, UK. [295]Cardiovascular and Metabolic Disease Signature Research Program, Duke-NUS Medical School, Singapore, Singapore. [296]Department of Public Health Solutions, National Institute for Health and Welfare, Helsinki, Finland. [297]Department of Public Health, University of Helsinki, Helsinki, Finland. [298]Saudi Diabetes Research Group, King Abdulaziz University, Jeddah, Saudi Arabia. [299]Department of Genomic Medicine and Environmental Toxicology, Instituto de Investigaciones Biomedicas, Universidad Nacional Autonoma de Mexico, Mexico City, Mexico. [300]Department of Public Health and Clinical Nutrition, University of Eastern Finland, Kuopio, Finland. [301]Department of Medicine, Internal Medicine, Lausanne University Hospital (CHUV), Lausanne, Switzerland. [302]Department of Public Health Sciences, Wake Forest School of Medicine, Winston-Salem, NC, USA. [303]Faculty of Medical Sciences, Newcastle University, Newcastle upon Tyne, UK. [304]Beijing Tongren Eye Center, Beijing Key Laboratory of Intraocular Tumor Diagnosis and Treatment, Beijing Ophthalmology & Visual Sciences Key Lab, Beijing Tongren Hospital, Capital Medical University, Beijing, China. [305]Department of Public Health, Faculty of Medicine, University of Kelaniya, Ragama, Sri Lanka. [306]Department of Research and Evaluation, Kaiser Permanente of Southern California, Pasadena, CA, USA. [307]Institute for Molecular Bioscience, The University of Queensland, St Lucia, Queensland, Australia. [308]Kurume University School of Medicine, Kurume, Japan. [309]TUM School of Medicine, Technical University of Munich and Klinikum Rechts der Isar, Munich, Germany. [310]Department of Pediatrics, Division of Endocrinology, Stanford School of Medicine, Stanford, CA, USA. [311]Wellcome Centre for Human Genetics, Nuffield Department of Medicine, University of Oxford, Oxford, UK. [312]Department of Medicine, Division of General Internal Medicine, Massachusetts General Hospital, Boston, MA, USA. [313]Department of Medicine, General Internal Medicine, Massachusetts General Hospital, Boston, MA, USA. [314]Department of Medicine, Diabetes Unit and Endocrine Unit, Massachusetts General Hospital, Boston, MA, USA. [315]Department of Human Genetics, University of Michigan, Ann Arbor, MI, USA. [316]Centre for Genetics and Genomics Versus Arthritis, Division of Musculoskeletal and Dermatological Sciences, The University of Manchester, Manchester, UK. [317]Centre for Musculoskeletal Research, Division of Musculoskeletal and Dermatological Sciences, The University of Manchester, Manchester, UK. [318]Department of Biostatistics, University of Liverpool, Liverpool, UK. [319]Present address: Genentech, South San Francisco, CA, USA. [320]These authors contributed equally: Ji Chen, Cassandra N. Spracklen, Gaëlle Marenne, Arushi Varshney, Laura J. Corbin. [321]These authors jointly supervised this work: Stephen C. J. Parker, Karen L. Mohlke, Claudia Langenberg, Eleanor Wheeler, Andrew P. Morris, Inês Barroso.
